# Utilizing tumor microenvironment microbial profiles and host gene expressions for survival subtyping in diverse types of cancers

**DOI:** 10.1101/2023.05.18.541268

**Authors:** Haohong Zhang, Mingyue Cheng, Lei Ji, Kang Ning

## Abstract

The tumor microbiome, a complex community of microbes found in tumors, has been found to be linked to cancer development, progression, and treatment outcome. However, it remains a bottleneck in distangling the relationship between the tumor microbiome and patient survival. In this study, we aimed to decode this complex relationship by developing ASD-cancer (autoencoder-based subtypes detector for cancer), a semi-supervised deep learning framework that could extract survival-related features from tumor microbiome and transcriptome data, and identify patients’ survival subtypes. By using samples from The Cancer Genome Atlas database, we identified two statistically distinct survival subtypes across all 20 types of cancer. Our framework provided improved risk-stratification (e.g., for Liver hepatocellular carcinoma, LIHC, log-rank test, *P* = 8.12E-6) compared to PCA (e.g., for LIHC, log-rank test, *P* = 0.87), predicted survival subtypes accurately, and identified biomarkers for survival subtypes. Additionally, we identified potential interactions between microbes and genes that may play roles in survival. For instance, in LIHC, *Arcobacter*, *Methylocella*, and *Isoptericola* may regulate host survival through interactions with host genes enriched in the HIF-1 signaling pathway, indicating these species as potential therapy targets. Further experiments on validation dataset have also supported these patterns. Collectively, ASD-cancer has enabled accurate survival subtyping and biomarker discovery, which could facilitate personalized treatment for a broad-spectrum types of cancers.

## Introduction

Cancer is a disease characterized by heterogeneous histopathological, genomic, and transcriptomic profiles within both the tumor and its microenvironment, which contribute to variations in response rates to therapy and patient outcomes [1]. The current clinical approaches to many types of cancer entail manual histopathological assessment, where tumor invasion, anaplasia, necrosis, and mitoses are used for grading and staging patients to guide therapeutic decision-making [2]. However, the subjective interpretation of histopathologic features has been demonstrated to suffer from substantial inter-and intraobserver variability, resulting in varying outcomes for patients with the same grade or stage. Therefore, identification of cancer subtypes is essential diagnosis and prognosis in clinics. Molecular signatures that enable cancer classification of cancer beyond conventional methods, such as stage, grade, or tissue of origin, are crucial as they provide patients the opportunities for personalized treatment strategies [3]. This approach is particularly useful for subtypes with similar molecular and pathway alterations because it enables the application of the same treatment modalities. Survival stratified patient subtypes are an example of a cancer subtype with prognostic significance, as which provide valuable insights into the molecular factors associated with survival [4].

Traditionally, cancer has been regarded as a disease originating from alterations in the genetic makeup of human beings [5, 6]. As such, there is a long history linking tumor gene expression to cancer outcomes [7-10]. However, cancer is a complex disease that involves not only the host but also the tumor microenvironment (TME), a complex ecosystem that surrounds and interacts with cancer cells. Gene expression data is subject specific limitations and noises [11]. Solely concentrating on host gene expression may neglect other molecular mechanisms in the TME. Therefore, integrating of multi-omics data has the potential to augment our comprehension of cancer [12, 13], and paves the way for precision medicine, which offers individualized diagnosis, prognosis, treatment and care [14, 15].

The tumor microbiome is a complex and diverse community of microbes that inhabit human tumors and adjacent tissue [16]. Poore et al. [17] recently developed a computational workflow that utilizes two orthogonal microbial detection pipelines to obtain high quality microbial abundances from high-throughput sequencing data of human tumors. Tumor microbiome has been shown to play a function in the development [18], progression [19], and response to treatment [20] of various types of cancer. The relationship between the tumor microbiome and patient survival is the subject of ongoing research, as some studies [21, 22] have suggested that the microbiome of a tumor may impact patient survival. *Malassezia globose*, for instance, has been linked to an increased risk of death in breast cancer patients [22]. The integration of tumor microbiomes and transcriptomes can provide a deeper comprehension of the interactions between microbes and host genes, thereby improving patient prognosis by enabling clinicians to devise tailored treatment strategies based on the unique characteristics of each patient’s cancer. This strategy can lead to more efficient and personalized treatment options, which may ultimately enhance patient outcomes. Consequently, the study of cancer subtypes, including those involving the tumor microbiome, represents an essential area of research that holds tremendous promise for improving cancer diagnosis, treatment, and ultimately patient survival. However, a lack of knowledge regarding the interaction between host genes and tumor microbiome has precluded us from using them for accurate patient survival analysis.

In this work, we investigated the relationship between host genes and microbes in the context of cancer survival by integrating host transcriptome and tumor microbiome data. We developed a deep learning-based framework called ASD-cancer (autoencoder-based subtypes detector for cancer), which is a semi-supervised deep learning framework based on autoencoder for the detection of cancer survival subtypes. We applied this framework on RNA-seq and tumor microbiome data from 20 types of cancer from The Cancer Genome Atlas (TCGA [23]) database and identified survival subtypes with high quality of patient risk stratification. Compared to the conventional PCA technique, the ASD-cancer framework demonstrated superior performance. In addition, we analyzed the distribution of clinical stages in our survival subtypes and discovered that several cancers had a similar stage distribution in both subtypes. We also found that important biomarkers for classifying survival subtypes are likely not sensitive to clinical stages. Furthermore, our analysis revealed that the high-risk group had more cancer-related pathways compared to the low-risk group, and we identified potential survival-related pathways of interaction between microbes and host genes.

## Results

### Ensemble deep learning-based survival subtype detection model using multi-omics data

We concentrated on 20 distinct types of cancer that are represented in the TCGA database. These types of cancer affect a wide range of organs and tissues throughout the body and 15 of them has clinical stage information defined by AJCC Cancer Staging System [2] **(Fig.1a, Supplementary Table 1)**. We obtained survival subtypes for these types of cancer using paired RNA-seq and tumor microbiome data from the TCGA database.

Deep learning [24] is a type of machine learning that employs neural networks that can automatically learn and extract data features without the need for manual feature engineering. It has been extensively utilized in several areas of cancer analysis, including diagnosis [25], prediction of drug response [26], and drug discovery [27]. In this study, we proposed ASD-cancer (autoencoder-based subtypes detector for cancer), a semi-supervised deep learning framework based on autoencoder, a type of neural network that is trained to reconstruct its input data. It composed of two components: an encoder that maps the input data to a lower-dimensional space, and a decoder that maps this representation back to the original data space. In our study, autoencoder models were used to extract relevant features from the normalized microbiome abundance data and transcriptome data for identifying cancer survival subtypes **(Fig. 1b)**. These extracted features were then analyzed using univariate Cox-PH regression to identify a subset of survival-related features. To ensure an adequate number of features, we implemented an ensemble step using a total of 20 models. We then determine the number of survival subtypes using Gaussian Mixture Models and the highest silhouette score (**Methods**).

**Fig. 1.**
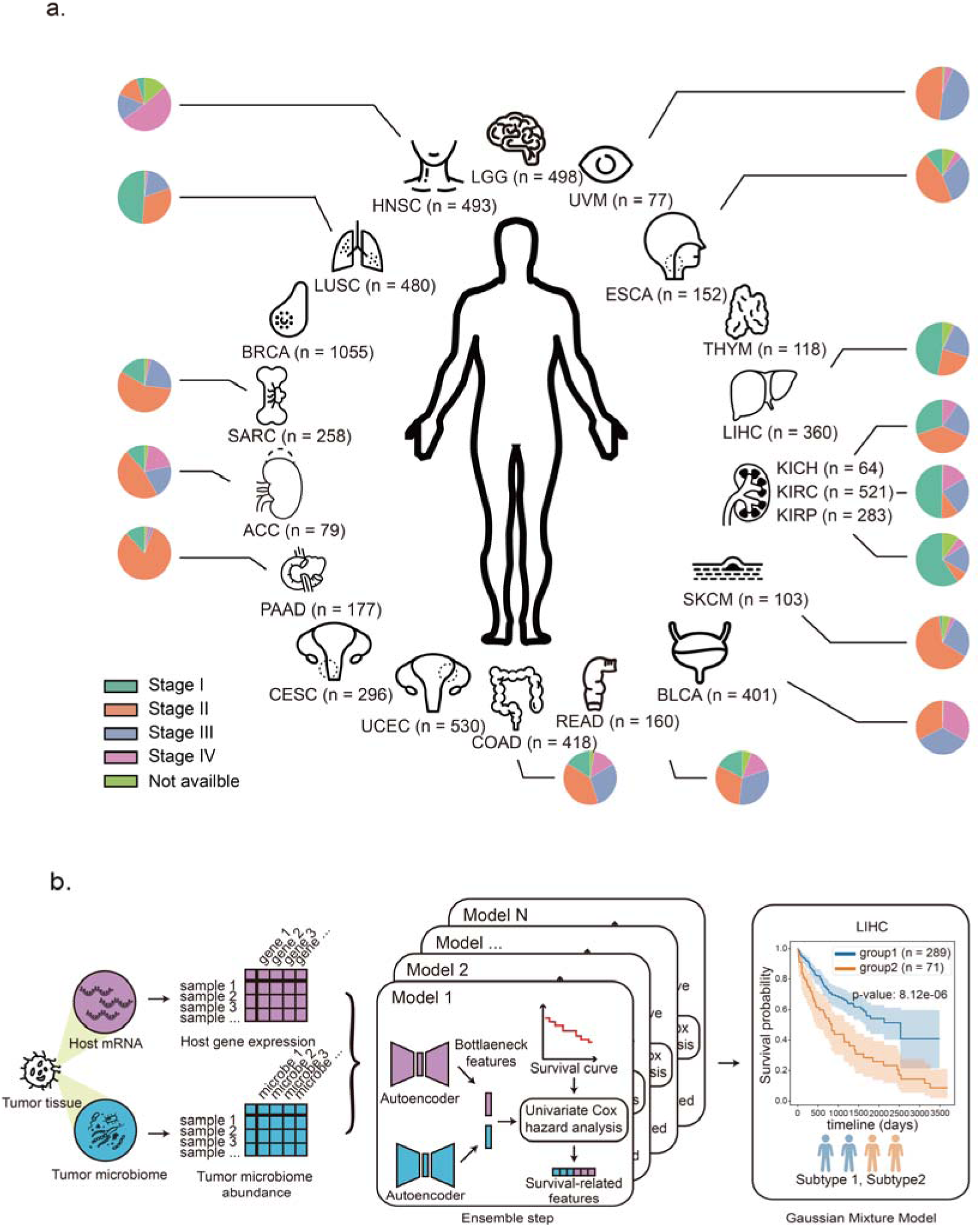
Material and pipeline of survival subtypes detection. **a.** We obtain the 20 cancer datasets from TCGA. Each dataset contains paired RNA-seq data and tumor microbiome data. The pie plot near each cancer represents the distribution of tumor stage defined by AJCC Cancer Staging System. The abbreviated names of cancer: ACC, Adrenocortical Carcinoma; BLCA, Bladder Urothelial Carcinoma; BRCA, Breast Invasive Carcinoma; CESC, Cervical Squamous Cell Carcinoma and Endocervical Adenocarcinoma; COAD, Colon Adenocarcinoma; ESCA, Esophageal Aarcinoma; HNSC, Head and Neck Squamous Cell Carcinoma; KICH, Kidney Chromophobe; KIRC, Kidney Renal Clear Cell Carcinoma; KIRP, Kidney Renal Papillary Cell Carcinoma; LGG, Brain Lower Grade Glioma; LIHC, Liver Hepatocellular Carcinoma; LUSC, Lung Squamous Cell Carcinoma; PAAD, Pancreatic Adenocarcinoma; READ, Rectum Adenocarcinoma; SARC, Sarcoma; SKCM, Skin Cutaneous Melanoma; THYM, Thymoma ; UCEC, Uterine Corpus Endometrial Carcinoma; UVM, Uveal Melanoma. The number of samples followed the abbreviated names. **b.** The pipeline of Ensemble deep learning-based survival subtype detection model.

### Two survival subtypes detected on 20 types of cancer

Using ASD-cancer, we analyzed the host RNA-seq and tumor microbiome data (**Methods**) of 20 TCGA cancers. For every type of cancer, we determined the optimal clustering number K that yields the best silhouette score, a metric that measures clustering stability and accuracy. Our analysis revealed that setting K to 2 yielded the highest silhouette score for each of the 20 types of cancer, thereby enabling the detection of two distinct survival subtypes **(Fig. 2a)**. Remarkably, the subtype with better prognosis outcomes was designated as ASD-1, while the subtype with worse prognosis outcomes as ASD-2. Using log-rank tests, we identified statistically significant differences (*P* < 0.05) between the two subtypes’ Kaplan-Meier curve. We also obtained high C-indexes (greater than 0.5, higher than the expected value of random models). In contrast to the utilization of a single omic, the integration of two omics demonstrated greater significance (**Supplementary Fig. 1**).

**Fig. 2.**
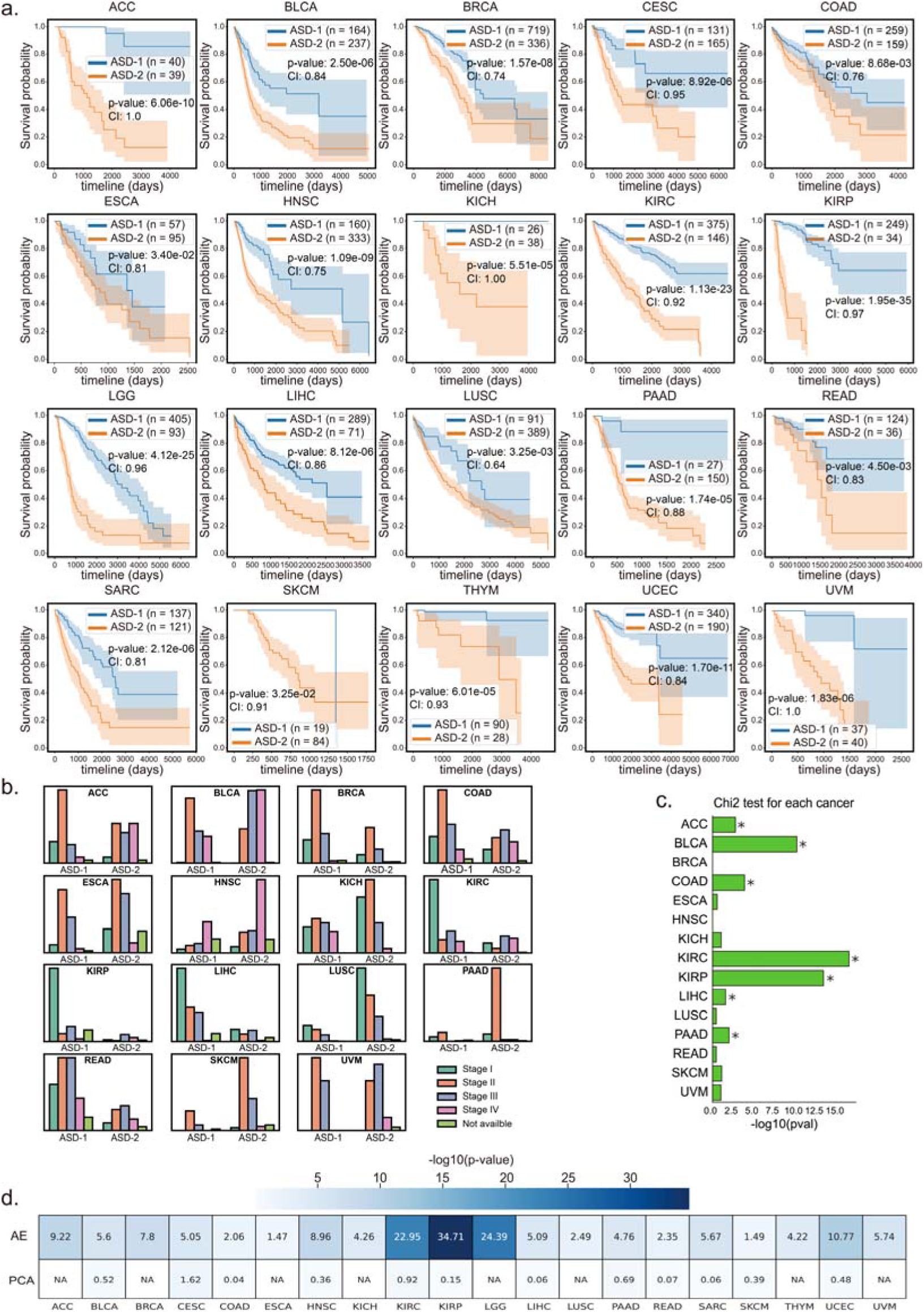
Subtypes detection for the 20 TCGA cancer datasets. **a.** Kaplan-Meier plots for each type of cancer. The survival curve for subtypes with better survival outcomes is marked in blue, while the curve for subtypes with worse survival outcomes is marked in orange. The p-value is the result of a log-rank test, which is a statistical test used to compare the survival curves of different groups. The CI is concordance index, a measurement of how well a model predicts the ordering of patients’ death times. **b.** Count plot showing the distribution of cancer stages among the subtypes. Five types of cancer (CESC, LGG, SARC, THYM, UCEC) do not have stage information. **c.** Bar plot showing the -log10 p value of a chi-squared test for the distribution of tumor stages among the two subtypes of the 16 types of cancer with stage information. A star symbol indicates a p-value less than 0.05. **d.** Heatmap showing the results of replacing the autoencoder with PCA. The values in the heatmap represent the -log10 of the p-value from a log-rank test. “NA” indicates that no survival-related features were extracted from at least one of the two omics data sets (RNA-seq and tumor microbiome) for a type of cancer. AE: autoencoder.

We also examined the distribution of age and clinical stage among the patients assigned to the two subtypes. Intriguingly, our analysis demonstrated that the age distributions of the two subtypes were identical across all types of cancer **(Supplementary Fig. 2)**. To investigate the relationship between the survival subtypes and clinical stages, chi-squared analyses were performed on the distribution of clinical stages between the two subtypes for each cancer. Our findings indicated that, among the 15 types of cancer with clinical stage information **(Fig. 2b, c, Supplementary Table 2)**, 7 of them, whose ASD-1 and ASD-2 could usually be differentiated clearly, exhibited significant differences (*P* < 0.05) in clinical stage distributions between the two subtypes. Among these seven types of cancer, ACC and BLCA exhibited overall survival-related cancer stage distributions, indicating that the survival subtyping of these cancers is highly related to clinical stage. Conversely, KIRC, KIRP, LIHC, PAAD, and COAD had a high proportion of a specific cancer stage in one subtype, indicating that the prognosis of these cancers is highly related to a specific clinical stage. For instance, advanced-stage KIRC patients were more likely to be stratified to the low-risk group. In contrast, for the remaining eight types of cancer whose ASD-1 and ASD-2 usually could not be differentiated clearly, we found no significant differences in the cancer stage distributions between the two subtypes. We propose that the prognostic outcome of these eight cancers may not be determined by clinical stages but by other molecular mechanisms.

We conducted an additional benchmark analysis by replacing the autoencoder module in ASD-cancer with a conventional Principal Component Analysis (PCA) decomposition **(Fig. 2d)**. To be specific, we transformed each omic dataset into 100 new components and followed the identical subsequent procedures as in the ASD-cancer study. Subtyping results showed that PCA failed to extract survival-related features in eight cancers, and 12 cancers had significantly poorer performances compared to the autoencoder module in ASD-cancer. These findings highlight the potential of deep learning and ensemble learning techniques, such as ASD-cancer, in extracting comprehensible features for the purpose of reliable cancer subtyping.

### Integrated multi-omics data shows high subtypes prediction accuracy

We conducted an analysis of the alpha diversity of tumor microbiomes in two subtypes of different types of cancer. The results revealed noteworthy variations in tumor microbiome alpha diversity between the two subtypes of nine types of cancer (ACC, BRCA, CESC, HNSC, LIHC, LUSC, LGG, KICH, and KIRP) **(Fig. 3a)**. Additionally, it was observed that the ASD-2 subtype exhibited elevated alpha diversity in their tumor microbiomes in comparison to the other five types of cancer (ACC, BRCA, HNSC, LIHC, LGG, and KIRP). We employed a Random Forest model to predict the survival subtypes of cancer **(Fig. 3b).** We selected the top 20 most important features from each omics (**Supplementary Table**) and developed models based on these features. The outcomes obtained from this approach were either consistent with or superior to those obtained from using all features. Finally, the integration of the two omics was performed and it was concluded that the model utilizing the chosen 40 features exhibited the highest level of precision. This model can be employed for prognosticating survival subtypes in forthcoming patients.

**Fig. 3.**
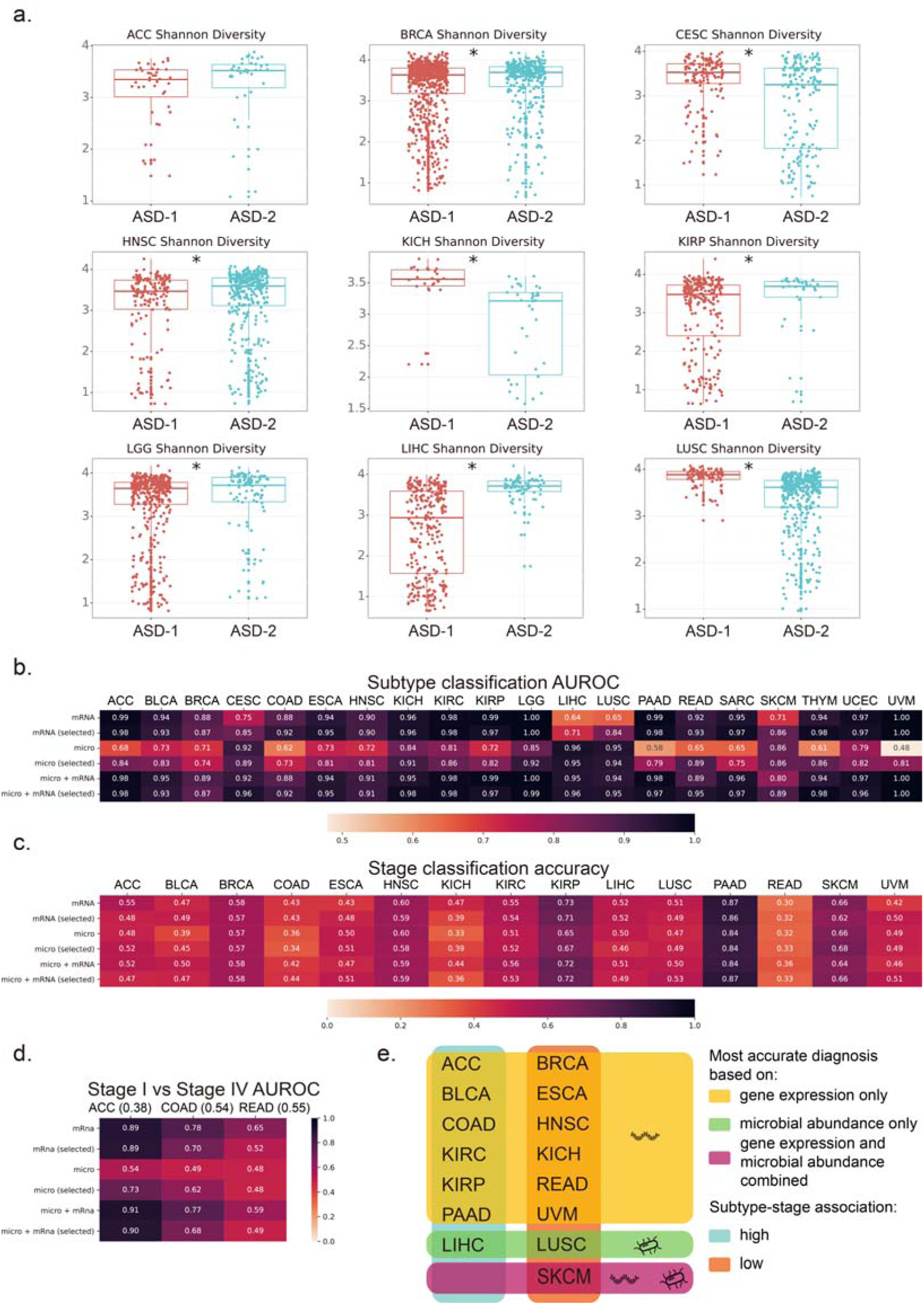
Alpha diversity of tumor microbiomes in different survival subtypes and the prediction results of subtype and stage. **a.** box plots of the significant differences in alpha diversity of tumor microbiomes between the two subtypes, with the test used being the Mann-Whitney test, and the star symbol indicating a p-value less than 0.05. **b.** the area under receiver operating characteristic (AUROC) heatmap of the prediction results of survival subtypes using a leave-one-out method in Random Forest. **c.** shows the accuracy heatmap of the prediction results of tumor stages using a leave-one-out method in Random Forest, with the numbers after each type of cancer indicating the number of stages. **d.** the accuracy heatmap of the prediction results of stage 1 and stage 4 using a leave-one-out method in Random Forest, with the ratios after each type of cancer indicating the proportion of stage I samples among the stage I and stage IV samples. In figures **b**, **c**, and **d**, each row represents different features. The first row represents all microbiome features; the second row represents the top 20 most important microbiome features; the third row represents all transcriptome features; the fourth row represents the top 20 most important transcriptome features; the fifth row represents all features from both omics’ approaches; the sixth row represents the top 20 most important features from both omics’ approaches. **e.** the categories of cancers based on the results of omics predictions. The first category consists of cancers with better results obtained from transcriptomics data and the results are consistent with transcriptomics when the two omics are integrated. The second category consists of cancers with better results obtained from microbiomes data and the results are consistent with microbiomes when the two omics are integrated. The third category consists of cancers with better results obtained from the integration of the two omics compared to using a single omics. Subtype-stage association: high means significant differences in clinical stage distributions between the two subtypes (chi-squared test, *P* <0.05), while low means no significant differences in clinical stage distributions between the two subtypes (chi-squared test, *P* > 0.05).

We also employed two omics to construct a Random Forest model for discriminating clinical stages among patients using the 40 features we selected above. Nonetheless, the attained accuracy levels for certain types of cancer were not deemed satisfactory **(Fig. 3c)**. Specifically, the accuracy for PAAD was 0.87, however, this outcome could be attributed to the substantial proportion of stage I patients present in the sample. Ultimately, our attention was directed towards the differentiation of patients in stage I and stage IV patients for types of cancer in which stage I or stage IV samples represented more than 30% of the total. We also excluded SKCM from our analysis due to the small number of samples. The final model was applied to three types of cancer: ACC, COAD, and READ **(Fig. 3d)**. The results indicate that the models for ACC and COAD performed well and that gene expression was more informative for distinguishing between stage I and stage IV than the tumor microbiome. These findings suggest that biomarkers used to discriminate survival subtypes may not be sensitive to clinical stages.

20 types of cancer were classified into three distinct categories on the basis of the predictive capabilities of transcriptomic and microbiome data. (**Fig. 3e**): best performance based on transcriptome only, microbiome only, and the integration of both omics. The first category exhibited superior transcriptome prediction results, the second category demonstrated better microbiome prediction results, and the third category showcased better outcomes from the integration of both omics. The results of our study indicate that the transcriptome is more strongly associated with predicting survival subtypes for the majority of cancers, with the exception of LIHC and LUSC, which exhibit a stronger correlation with the microbiome. The integration of both omics led to an enhanced prediction accuracy for CESC and SKCM, indicating a potential involvement of the interplay between tumor microbiomes and host genes in the survival of these two types of cancers.

### Multi-omics features that play critical roles in the **association of pre-defined cancer stages and intrinsic survival subtypes**

In this study, we ascertain the molecular mechanisms that underlie the variations in survival outcomes observed among different cancer subtypes. We utilized Gene Set Enrichment Analysis (GSEA) to compare the association of pre-defined cancer stages and intrinsic survival subtypes. Our results indicate that the ASD-2 subtype exhibited a higher degree of enrichment in cancer-related pathways as compared to the ASD-1 subtype in cancers whose clinical stage is related to survival subtyping.

Firstly, we analyzed BLCA and LIHC, whose survival-specific subtyping aligned with clinical stages. The result revealed that the ASD-2 subtype in BLCA demonstrated an increase in the Proteoglycans in Cancer and Relaxin signaling pathways (**Fig. 4a**), which are responsible for the regulation of cellular behavior, facilitation of tumor proliferation, and promotion of angiogenesis. We also observed that certain microbes such as *Shuttleworthia*, *Zymobacter*, and *Salinimicrobium* may interact with host genes like FGF2 and further modulate signaling pathways like MAPK and Ras in the ASD-2 subtype [28]. Similarly, in LIHC **(Fig. 4b)**, we found microbes such as Arcobacter, *Methylocella*, and *Isoptericola* also play a role in regulating host survival through interactions with host genes and regulation of critical signaling pathways and ASD-2 enrichment in the HIF-1 and Sphingolipid signaling pathways, which are involved in tumor angiogenesis and cancer development, respectively [29, 30].

**Fig. 4.**
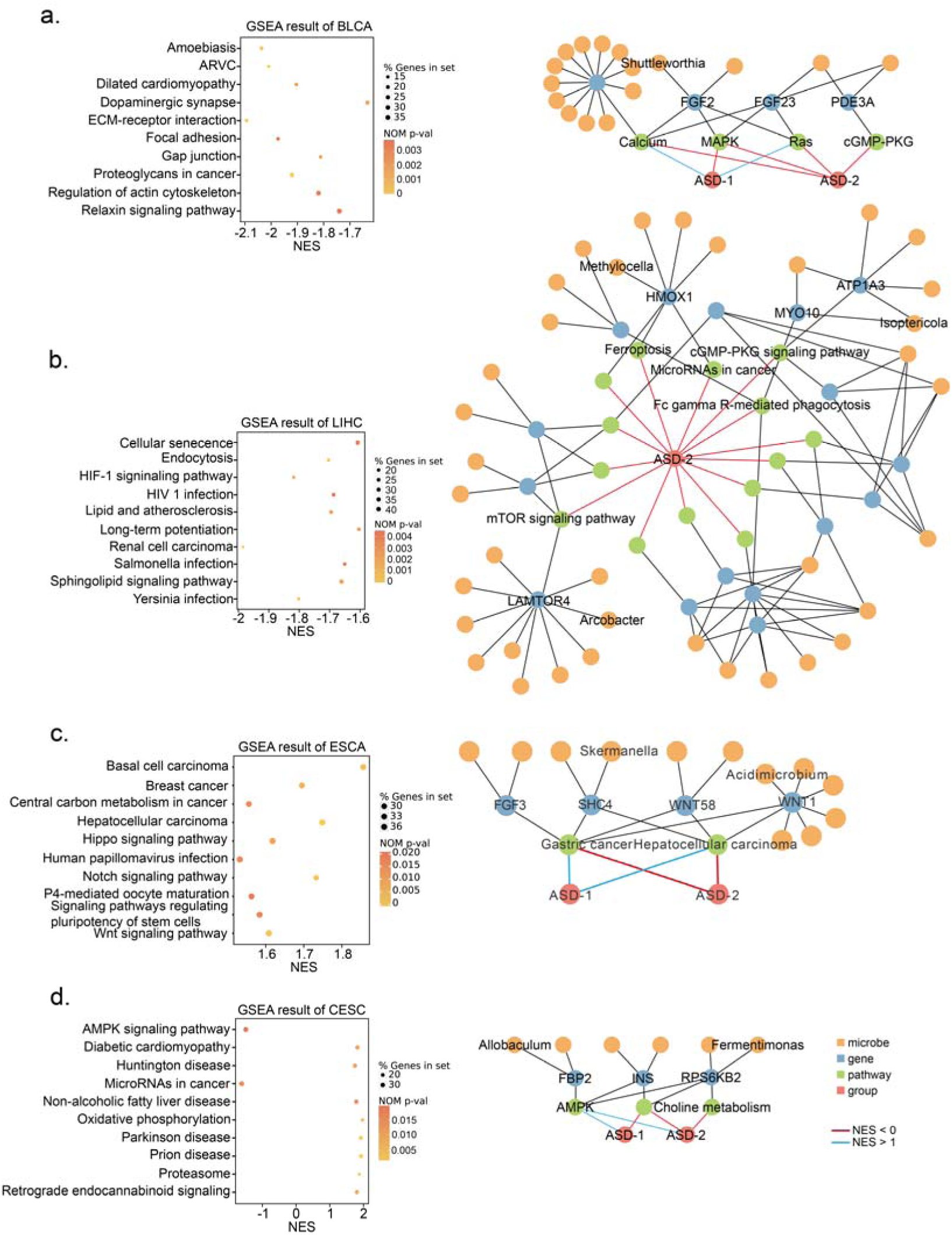
GSEA result and correlation network for three representative cancers. **a.** The tops 10 enriched pathways for BLCA based on Gene Set Enrichment Analysis (GSEA) and the correlation network between microbes and host genes (details see **Method**). **b.** The tops 10 enriched pathways for LIHC based on Gene Set Enrichment Analysis (GSEA) and the correlation network between microbes and host genes (details see **Method**). c. The top 10 enriched pathways for ESCA based on Gene Set Enrichment Analysis (GSEA) and the correlation network between microbes and host genes (details see **Method**). **d.** The tops 10 enriched pathways for CESC based on Gene Set Enrichment Analysis (GSEA) and the correlation network between microbes and host genes (details see **Method**). The depth of the point color represents the p-value of enrichment, and the size of the point represents the number of genes enriched in the pathway. The value on the x-axis is the enrichment score, with positive values representing enrichment in subtypes with better survival, and negative values representing enrichment in subtypes with worse survival.

We then analyzed ESCA, which had no significant difference in clinical stages between our two subtypes **(Fig. 4c)**. We found that the ASD-1 subtype was enriched in pathways not directly related to ESCA. We also found that for HNSC, BRCA, and LUSC, whose survival-specific subtyping did not align with clinical stages, there were no significant correlations between the microbiome and host genes. The implication of this statement is that the prognosis of these cancers is primarily determined by the distinct influence of omics rather than the interplay among multiple omics. Interestingly, The ASD-1 subtype of ESCA exhibited a notable enrichment of gastric cancer and hepatocellular carcinoma, both of which are classified as digestive system cancers.

Finally, we analyzed cancers without clinical stage information, such as CESC **(Fig. 4d)**. The ASD-2 subtype in CESC showed enrichment in pathways such as the AMPK signaling pathway and MicroRNAs in cancer. The promotion of tumor growth and chemotherapy resistance can be attributed to the dysregulation of the AMPK signaling pathway [31], whereas the dysregulation of miRNA expression is a prevalent characteristic of numerous types of cancer [32]. Furthermore, we found that certain microbes such as *Allobaculum* and *Fermentimonas* may interact with host genes like RPS6KB2 and regulate AMPK signaling and choline metabolism in cancer pathways [33].

In summary, this work offers deeper understanding regarding plausible omics patterns that may account for variations in survival results. The results of our study indicate that tumors exhibiting increased interaction between the microbiome of the tumor and the host gene are more strongly correlated with clinical stage and prognosis, whereas other tumors are more closely linked to individual omics. Additional investigation is required to authenticate these discoveries and clarify the function of microbes in controlling host viability and pathways associated with cancer.

### Validation of ASD-cancer performance by other cohorts

The ASD-cancer workflow demonstrates the capability to predict the survival subtype of new individual samples based on shared tumor microbiome and gene expression features. To further validate the patient survival risk stratification achieved by the ASD-cancer models, we conducted tests on additional independent cancer datasets obtained from the LIHC and PAAD cohorts within the International Cancer Genome Consortium (ICGC) database. The tumor microbiome abundance data was annotated using Poore et al.’s pipeline [17], and our trained model was applied to these two cohorts (**Methods**). The results yielded a C-index of XX and a log-rank p-value of XX for the LIHC cohort, and a C-index of XX and a log-rank p-value of XX for the PAAD cohort (**Fig. XX**). Furthermore, we developed a Random Forest model based on 40 features selected from our original Random Forest model. Notably, this Random Forest model achieved an AUROC of XX, demonstrating accurate and efficient prediction without the need for additional clinical variables or lower-dimensional feature transformations via the autoencoder method (**Fig. XX**). Consequently, we have successfully validated the predictability of ASD-cancer using the additional LIHC and PAAD cohorts.

## Discussion

Although the relationship between survival rates and clinical stage classification in cancer patients is controversial, it is generally accepted that, under optimal conditions, there is an inverse correlation between survival outcomes and clinical stage progression (**Fig. 5a**). However, patients with the same grade or stage demonstrate significant variability in outcomes. Although recent methods for cancer survival subtyping based on molecular signatures have emerged, they typically only focus on a single omic [11] or the host itself [34], and tend to ignore other complex factors in the TME, such as the tumor microbiome [12]. Our study found that survival-specific subtyping, by integrating tumor microbiome data and host gene expression data, does not fully align with clinical stage data. Seven types of cancer in our study showed high clinical stage and survival-subtyping association, whereas eight types showed no difference in clinical stage distribution between two survival subtypes (**Fig. 5b**). Our findings suggest that the tumor microbiome and host gene interact more actively in tumors with a high correlation between pre-defined cancer stages and intrinsic survival subtypes. Conversely, cancers with a low association between pre-defined cancer stages and intrinsic survival subtypes exhibit fewer interactivities between the tumor microbiome and host gene (**Fig. 5c**). These patterns indicate the two groups of cancer, depending on the interaction between host genes and tumor microbiomes: In the first group, host genes and tumor microbiomes has a weak correlation, and ASD-1 and ASD-2 usually could not be differentiated clearly. For example, ESCA in first group has a log-rank test p-value 3.40e-02. A more concrete example is on LUSC, for which we found no strong correlation pathway between tumor microbiome and host gene (R^2^ < 0.9), implying that the interactivity between tumor microbiome and host genes in these cancers is weak, but the prognosis of these cancers is primarily determined by the distinct influence of each omic, which is responsible for survival-specific subtyping in this particular cancer (log-rank test p-value 3.25e-03). While in the second group, host genes and tumor microbiomes have strong correlation, and ASD-1 and ASD-2 usually could be differentiated clearly. For example, LIHC in second group has log-rank test p-value 8.12e-06. Collectively, these patterns have not only demonstrated that microbes in the tumor microenvironment might influence cancer development in a variety of ways, but also that we can subtype patients with high accuracy based on their prognosis using tumor microbiome and host gene expressions, which is a critical addition to clinical stage information for monitoring cancer patients’ status.

**Fig. 5.**
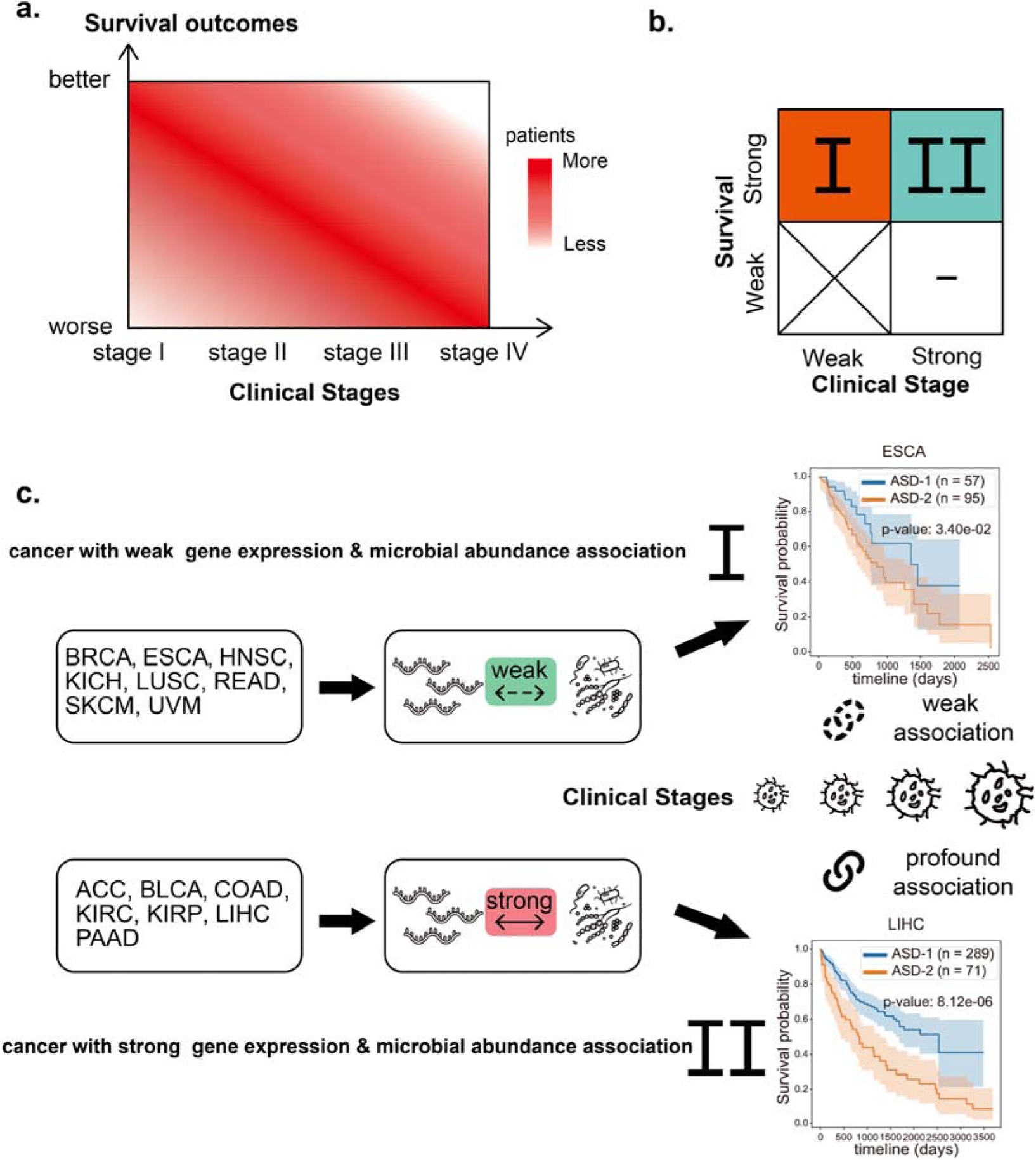
Illustration of Clinical Stages of Tumor and Survival Subtyping. **a.** Heatmap of the distribution of patients’ survival rates across different clinical stages of tumors under the ideal state. The x-axis represents the clinical stages of the tumor, and the y-axis represents the survival outcomes. **b.** Two groups of cancer identified in our study. In the first group, there is weak association between clinical stages and survival subtypes, while in the second group, there is strong association between clinical stages and survival subtypes. **c.** Schematic representation of the interaction between host genes and tumor microbiomes in the two groups of cancer. In the first group, host genes and tumor microbiomes are weakly correlated, and ASD-1 and ASD-2 usually indistinguishable; while in the second group, host genes and tumor microbiomes are strongly correlated, and ASD-1 and ASD-2 are typically distinguishable.

Furthermore, we found that compared to the low-risk group, the high-risk group was more likely to have a greater number of pathways connected to cancer. This provides evidence that poor survival outcomes in cancer patients may be driven by distinct biological pathways. In addition, we found numerous possible microbial-gene interaction pathways that might contribute to cancer survival. For example, in the case of LIHC, *Arcobacter*, *Methylocella*, and *Isoptericola* may regulate host survival through interactions with host genes enriched in critical signaling pathways in cancer, particularly the HIF-1 signaling pathway. These species could be targeted for therapy to improve patient prognosis. This discovery is significant because it implies the tumor microbiome may play a pivotal role in determining cancer patients’ chances of survival.

In conclusion, our study supports the idea that integrating transcriptome and microbiome data can aid in understanding the factors that influence cancer patients’ survival. Specifically, we focused on ASD-cancer among 20 types of cancer and discovered two survival subtypes, ASD-1 and ASD-2, which are not solely reliant on traditional clinical criteria such as cancer stage and patient age. Additionally, we successfully categorized 15 out of the 20 types of cancer with clinical stage information into two groups, based on the relationship between pre-defined cancer stages and intrinsic survival subtypes. This revealed different interactive patterns between host genes and tumor microbiome. These findings have the potential to enhance the prognostic information accessible to clinicians and contribute to the developing field of precision medicine. Our study sheds light on the potential for using multi-omics data and deep learning method to improve cancer prognosis and discover personalized therapeutic targets.

## Methods

### TCGA datasets

Transcriptome data for this study was obtained from the TCGA database (https://brd.nci.nih.gov/brd/sop-compendium/show/701). Only SampleIDs containing ‘01A’ were retained to ensure the data came from primary tumors. The tumor microbiome data was obtained from a previous cancer microbiome study [17], which included whole genome and whole transcriptome sequencing data of 33 types of cancers from the TCGA database. The data was processed using state-of-the-art tools to minimize sample contamination [35], and only data derived from whole transcriptome sequencing was used. The final result was a paired tumor microbiome and transcriptome dataset comprising 20 types of cancer from tissue samples.

### ASD-cancer workflow

ASD-cancer workflow was designed in several modules as follow: The first module preprocess the data, which involves normalization and scaling of both mRNA and microbiome data to ensure comparability between the two datasets. The second module transformed each omic feature into low-demension representation using an autoencoder. An ensemble step was used to obtain adequate features by training 20 autoencoders for each cancer. The third module establishes cancer survival subtypes by performing a univariate Cox proportional hazards analysis and Gaussian mixture model clustering. The fourth module establishes the association of cancer survival subtypes and stages using chi-square tests and random forest models for subtype and stage prediction. The fifth module identifies the association between host gene expressions and microbial abundances using Pearson correlation analysis. Finally, gene enrichment analysis was performed to identify biological pathways associated with different cancer survival subtypes.

### Preprocessing

During the preprocessing stage, normalization was performed on both mRNA and microbiome data to ensure comparability between the two datasets for downstream analyses. The expression values of the mRNA data were normalized using the RankNorm method, while the microbiome data was normalized by dividing each value in a row by the sum of all values in that row. After normalization, both datasets were further scaled using the StandardScaler method from the Scikit-learn (sklearn) library. The normalization and scaling procedures are essential in addressing the differences in the magnitudes and distributions of the data, thus enabling reliable comparisons and joint analysis of the datasets.

### Autoencoder transformation

Autoencoders are a type of neural network used for unsupervised learning that can be used to learn efficient data representations in an unsupervised way. The network consists of two main components: the encoder, which maps the input data to a lower-dimensional representation, and the decoder, which reconstructs the original data from the encoded representation. The general formula for an autoencoder can be written as follows:

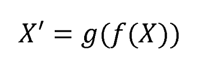

where X is the input data, f is the encoder function, g is the decoder function, and X’ is the reconstructed output.

In this study, we utilized an autoencoder to transform each omic data into a lower-dimensional representation. We implemented an ensemble step to obtain adequate features by training 20 autoencoders for each cancer. Each autoencoder had an encoder with two linear layers of size 500 and 100 respectively, with a dropout rate of 0.2 and Tanh activation function in between. The decoder also had two linear layers of size 500 and the length of each omic feature respectively with a dropout of 0.2 and Tanh activation function in between. We used the PyTorch package to implement the model. In the end, we obtained 100*20 lower-dimensional features for each cancer.

### Cancer survival subtyping

Cancer survival subtyping was established by performing a univariate Cox proportional hazards (Cox-PH) analysis on 100*20 features that had been reduced in dimension using autoencoder, along with survival information for the samples. We identified the features that were significantly associated with survival by selecting those with p-values < 0.05. Using all the selected features, we performed Gaussian mixture model (GMM) clustering with K values ranging from 2-5. We compared the silhouette score for each cluster, and selected the K value that resulted in the highest silhouette score as the number of clusters. Once we obtained the clustering results, we plotted Kaplan-Meier curves for each cluster and performed the log-rank test to assess the differences in survival between the clusters. We used the lifelines package for Cox-PH analysis, Kaplan-Meier curve plotting, and log-rank test, and the scikit-learn package for GMM clustering.

### Establishment of the association of cancer survival subtypes and stages

We first analyzed the distribution of clinical stage across different cancer subtypes and used a chi-square test to evaluate whether there were significant differences in clinical stage distribution between the subtypes. Next, we used a random forest model to predict the cancer subtype and clinical stage for each sample. For subtype prediction, we used leave-one-out cross-validation to calculate the area under the receiver operating characteristic curve (AUROC) and selected the top 20 features based on their importance scores. For clinical stage prediction, we calculated the accuracy using leave-one-out cross-validation. We selected cancers for stage I and stage IV prediction based on the proportion of samples in stage I or stage IV was at least 30% and excluded skin cutaneous melanoma (SKCM) due to a small sample size.

### Establishment of the association of host gene expressions and microbial abundances

To explore the relationship between host gene expressions and microbial abundances, we first calculated the correlation coefficients between each host gene and microbial species using Pearson correlation analysis. Specifically, we utilized PyTorch to implement the correlation analysis and utilized GPU acceleration to speed up the computation. We only selected microbiome-host gene pairs with a correlation coefficient greater than 0.9 to ensure high confidence in the correlation.

To identify the biological pathways associated with different cancer survival subtypes, we performed gene enrichment analysis using the gseapy package. This package provides access to various databases and tools for gene set enrichment analysis, including the KEGG and Reactome databases. We calculated the enrichment scores for each pathway and used the q-value to select the top 10 enriched pathways for visualization.

### Visualization of results

We utilized several software tools to visualize the results of our analyses. Cytoscape was used to create the microbiome-host gene correlation network. Kaplan-Meier curves were plotted using the lifelines package to compare the survival of different subtypes. For other types of visualizations such as AUROC and accuracy heatmaps, we utilized the plotnine and seaborn packages.

### Statistical analysis

All statistical analyses were performed using Python packages, including Scikit-learn, scipy, gseapy, and lifelines. The significance level was set at 0.05, and all p-values were two-sided.

### Code availability

All source code has been uploaded to the website at https://github.com/HUST-NingKang-Lab/ASD-cancer

## Acknowledgments

This work was partially supported by the National Natural Science Foundation of China (Grant Nos. 32071465, 31871334, and 31671374), and the National Key R&D Program of China (Grant No. 2018YFC0910502). Numerical computations were performed on the Hefei Advanced Computing Center.

## Ethics approval and consent to participate

Not applicable.

