## Supplementary Fig. 1, Supplementary Fig. 2, Supplementary Fig. 3, Supplementary Table 1, Supplementary Table 2, Supplementary Table 3 for "Utilizing tumor microenvironment microbial profiles and host gene expressions for survival subtyping in diverse types of cancers"

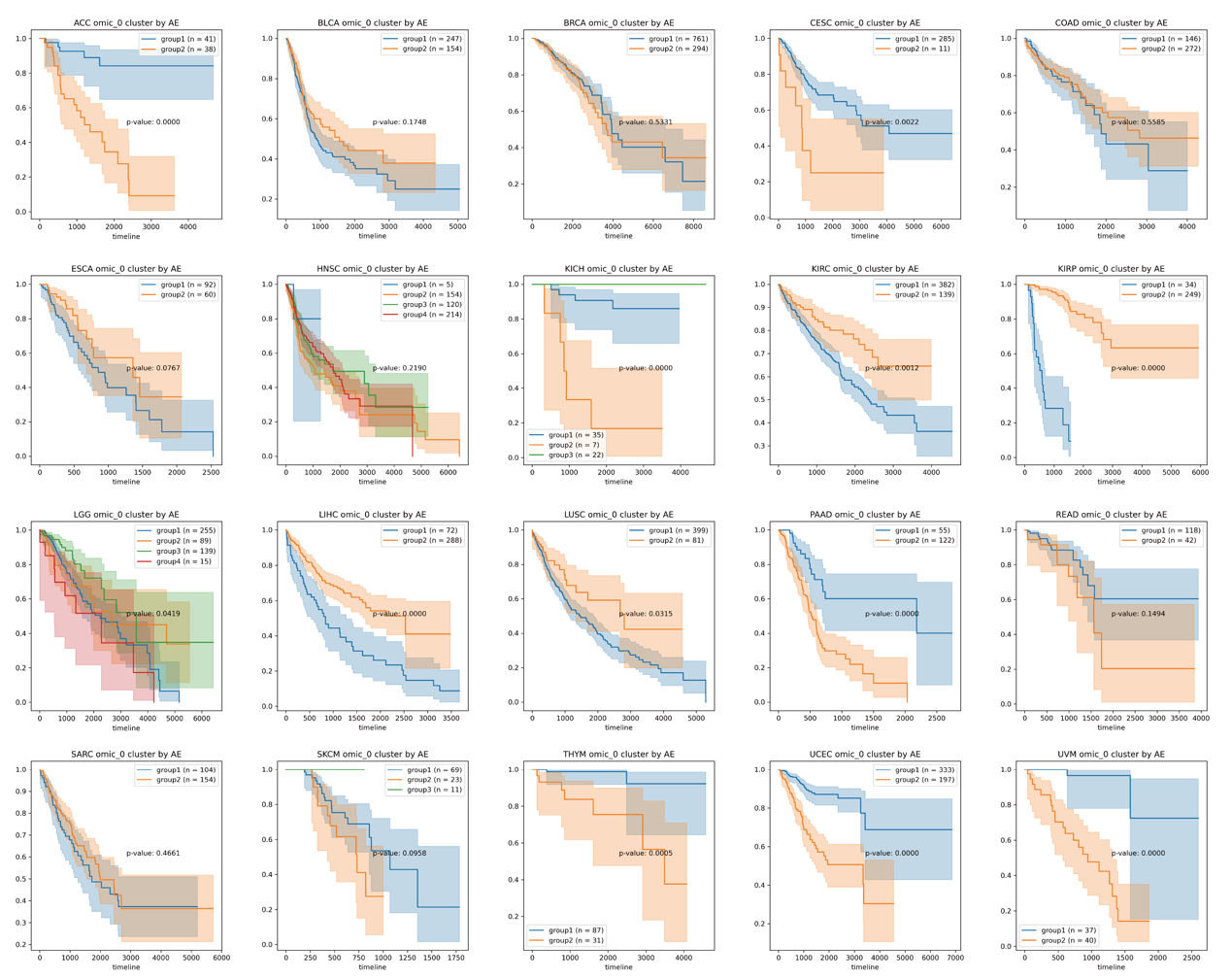

**Supplementary Figure 1. Subtyping result only using transcriptome.** Kaplan-Meier plots for each type of cancer. The p-value is the result of a log-rank test, which is a statistical test used to compare the survival curves of different groups.

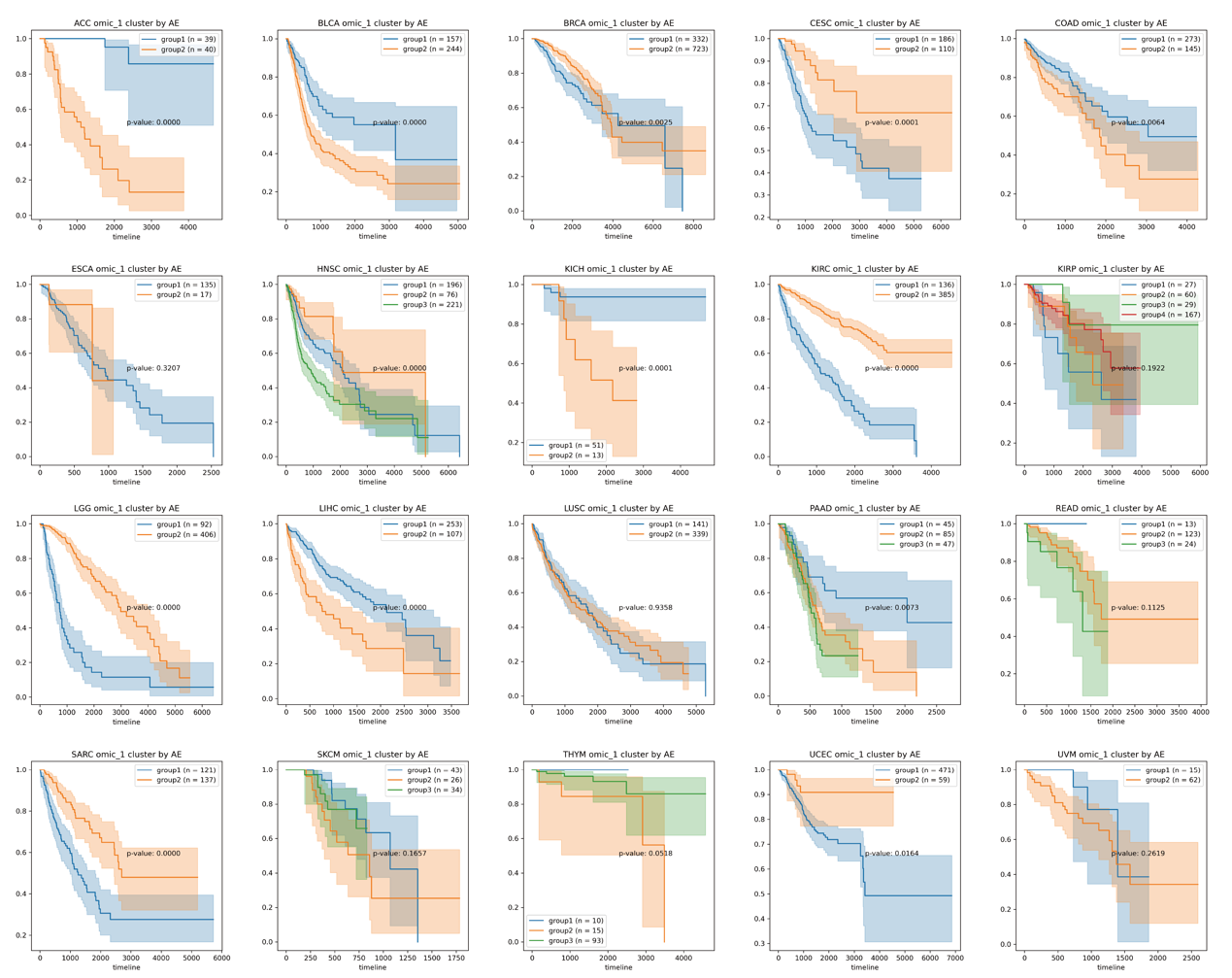

**Supplementary Figure 2. Subtyping result only using tumor microbiome.** Kaplan-Meier plots for each type of cancer. The p-value is the result of a log-rank test, which is a statistical test used to compare the survival curves of different groups.

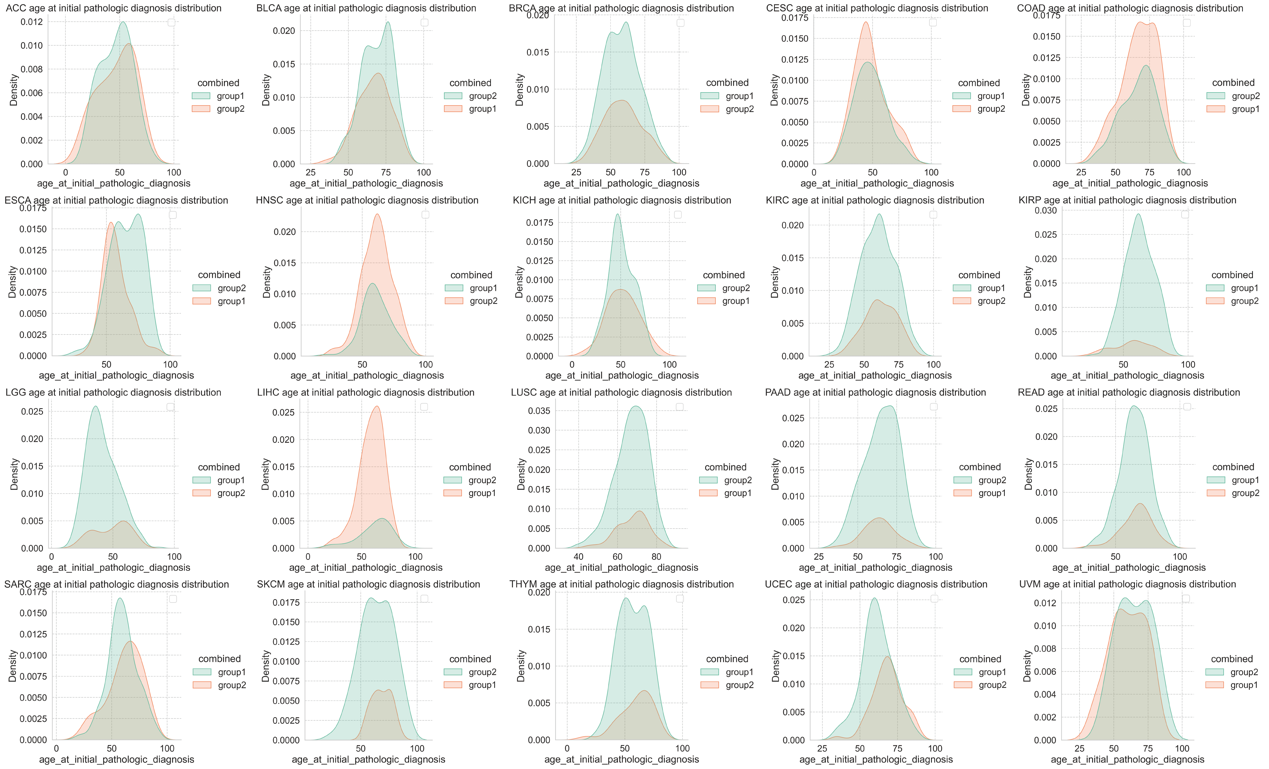

**Supplementary Figure 3. Age distribution in ASD-1 and ASD-2 among 20 types of cancer.**

| **Cancer** | **Stage I** | **Stage II** | **Stage III** | **Stage IV** | **Not available** | **Total** |
| --- | --- | --- | --- | --- | --- | --- |
| ACC | 9 | 37 | 16 | 15 | 2 | 79 |
| BLCA | 1 | 130 | 137 | 131 | 2 | 401 |
| BRCA | 174 | 598 | 240 | 20 | 23 | 1055 |
| CESC | 0 | 0 | 0 | 0 | 296 | 296 |
| COAD | 67 | 163 | 119 | 58 | 11 | 418 |
| ESCA | 16 | 69 | 48 | 7 | 12 | 152 |
| HNSC | 24 | 69 | 78 | 255 | 67 | 493 |
| KICH | 19 | 25 | 14 | 6 | 0 | 64 |
| KIRC | 259 | 55 | 122 | 82 | 3 | 521 |
| KIRP | 169 | 20 | 50 | 15 | 29 | 283 |
| LGG | 0 | 0 | 0 | 0 | 498 | 498 |
| LIHC | 168 | 84 | 80 | 6 | 22 | 360 |
| LUSC | 234 | 152 | 83 | 7 | 4 | 480 |
| PAAD | 21 | 146 | 3 | 4 | 3 | 177 |
| READ | 28 | 49 | 51 | 23 | 9 | 160 |
| SARC | 0 | 0 | 0 | 0 | 258 | 258 |
| SKCM | 2 | 66 | 27 | 3 | 5 | 103 |
| THYM | 0 | 0 | 0 | 0 | 118 | 118 |
| UCEC | 0 | 0 | 0 | 0 | 530 | 530 |
| UVM | 0 | 37 | 35 | 4 | 1 | 77 |

**Supplementary Table 1. Sample counts and stage distribution of each cancer.** ACC, Adrenocortical Carcinoma; BLCA, Bladder Urothelial Carcinoma; BRCA, Breast Invasive Carcinoma; CESC, Cervical Squamous Cell Carcinoma and Endocervical Adenocarcinoma; COAD, Colon Adenocarcinoma; ESCA, Esophageal Aarcinoma; HNSC, Head and Neck Squamous Cell Carcinoma; KICH, Kidney Chromophobe; KIRC, Kidney Renal Clear Cell Carcinoma; KIRP, Kidney Renal Papillary Cell Carcinoma; LGG, Brain Lower Grade Glioma; LIHC, Liver Hepatocellular Carcinoma; LUSC, Lung Squamous Cell Carcinoma; PAAD, Pancreatic Adenocarcinoma; READ, Rectum Adenocarcinoma; SARC, Sarcoma; SKCM, Skin Cutaneous Melanoma; THYM, Thymoma ; UCEC, Uterine Corpus Endometrial Carcinoma; UVM, Uveal Melanoma

**Supplementary Table 2. Top 20 outstanding microbial species with highest contribution of each cancer in subtype prediction.**

| **Adrenocortical Carcinoma** | | |
| --- | --- | --- |
| **Ranking** | **Microbial Types** | **Contribution (×10^­-3^)** |
| 1 | k__Bacteria.p__Proteobacteria.c__Gammaproteobacteria.o__Thiotrichales.f__Francisellaceae.g__Francisella | 13.75 |
| 2 | k__Bacteria.p__Firmicutes.c__Bacilli.o__Bacillales.f__Bacillaceae.g__Domibacillus | 12.34 |
| 3 | k__Bacteria.p__Firmicutes.c__Clostridia.o__Thermoanaerobacterales.f__Thermoanaerobacterales_Family_III._Incertae_Sedis.g__Caldicellulosiruptor | 10.51 |
| 4 | k__Bacteria.p__Proteobacteria.c__Gammaproteobacteria.o__Enterobacteriales.f__Enterobacteriaceae.g__Citrobacter | 9.81 |
| 5 | k__Bacteria.p__Proteobacteria.c__Alphaproteobacteria.o__Rhizobiales.f__Methylocystaceae.g__Methylocystis | 9.62 |
| 6 | k__Bacteria.p__Proteobacteria.c__Gammaproteobacteria.o__Enterobacteriales.f__Enterobacteriaceae.g__Proteus | 9.10 |
| 7 | k__Bacteria.p__Proteobacteria.c__Gammaproteobacteria.o__Enterobacteriales.f__Enterobacteriaceae.g__Escherichia | 8.53 |
| 8 | k__Bacteria.p__Bacteroidetes.c__Bacteroidia.o__Bacteroidales.f__Bacteroidaceae.g__Bacteroides | 8.31 |
| 9 | k__Bacteria.p__Bacteroidetes.c__Flavobacteriia.o__Flavobacteriales.f__Flavobacteriaceae.g__Xanthomarina | 7.74 |
| 10 | k__Bacteria.p__Firmicutes.c__Bacilli.o__Lactobacillales.f__Streptococcaceae.g__Streptococcus | 7.33 |
| 11 | k__Viruses.o__Herpesvirales.f__Herpesviridae.g__Simplexvirus | 7.15 |
| 12 | k__Bacteria.p__Firmicutes.c__Clostridia.o__Clostridiales.f__Ruminococcaceae.g__Acetivibrio | 6.99 |
| 13 | k__Bacteria.p__Bacteroidetes.c__Flavobacteriia.o__Flavobacteriales.f__Flavobacteriaceae.g__Algibacter | 6.80 |
| 14 | k__Bacteria.p__Firmicutes.c__Bacilli.o__Lactobacillales.f__Streptococcaceae.g__Lactococcus | 6.69 |
| 15 | k__Bacteria.p__Proteobacteria.c__Alphaproteobacteria.o__Rhizobiales.f__Bartonellaceae.g__Bartonella | 6.51 |
| 16 | k__Bacteria.p__Firmicutes.c__Bacilli.o__Lactobacillales.f__Lactobacillaceae.g__Lactobacillus | 6.29 |
| 17 | k__Bacteria.p__Fusobacteria.c__Fusobacteriia.o__Fusobacteriales.f__Leptotrichiaceae.g__Sneathia | 6.13 |
| 18 | k__Bacteria.p__Proteobacteria.c__Deltaproteobacteria.o__Desulfobacterales.f__Desulfobulbaceae.g__Desulfotalea | 6.09 |
| 19 | k__Bacteria.p__Proteobacteria.c__Gammaproteobacteria.o__Enterobacteriales.f__Enterobacteriaceae.g__Trabulsiella | 5.95 |
| 20 | k__Bacteria.p__Firmicutes.c__Bacilli.o__Lactobacillales.f__Aerococcaceae.g__Aerococcus | 5.82 |

| **Bladder Urothelial Carcinoma** | | |
| --- | --- | --- |
| **Ranking** | **Microbial Types** | **Contribution (×10^­-3^)** |
| 1 | k__Bacteria.p__Firmicutes.c__Tissierellia.o__Tissierellales.f__Peptoniphilaceae.g__Parvimonas | 14.49 |
| 2 | k__Bacteria.p__Proteobacteria.c__Betaproteobacteria.o__Burkholderiales.f__Oxalobacteraceae.g__Collimonas | 9.52 |
| 3 | k__Viruses.f__Nimaviridae.g__Whispovirus | 7.52 |
| 4 | k__Bacteria.p__Proteobacteria.c__Betaproteobacteria.o__Burkholderiales.f__Sutterellaceae.g__Sutterella | 7.11 |
| 5 | k__Bacteria.p__Proteobacteria.c__Gammaproteobacteria.o__Pasteurellales.f__Pasteurellaceae.g__Gallibacterium | 6.89 |
| 6 | k__Bacteria.p__Bacteroidetes.c__Flavobacteriia.o__Flavobacteriales.f__Flavobacteriaceae.g__Apibacter | 5.91 |
| 7 | k__Bacteria.p__Proteobacteria.c__Alphaproteobacteria.o__Rhizobiales.f__Hyphomicrobiaceae.g__Prosthecomicrobium | 5.14 |
| 8 | k__Bacteria.p__Proteobacteria.c__Alphaproteobacteria.o__Rhodobacterales.f__Rhodobacteraceae.g__Paracoccus | 5.00 |
| 9 | k__Archaea.p__Euryarchaeota.c__Halobacteria.o__Haloferacales.f__Haloferacaceae.g__Haloquadratum | 4.53 |
| 10 | k__Bacteria.p__Proteobacteria.c__Gammaproteobacteria.o__Alteromonadales.f__Psychromonadaceae.g__Psychromonas | 4.50 |
| 11 | k__Bacteria.p__Bacteroidetes.c__Cytophagia.o__Cytophagales.f__Cytophagaceae.g__Anditalea | 4.34 |
| 12 | k__Bacteria.p__Actinobacteria.c__Actinobacteria.o__Pseudonocardiales.f__Pseudonocardiaceae.g__Amycolatopsis | 4.07 |
| 13 | k__Bacteria.p__Actinobacteria.c__Actinobacteria.o__Micrococcales.f__Intrasporangiaceae.g__Terrabacter | 3.99 |
| 14 | k__Bacteria.p__Proteobacteria.c__Alphaproteobacteria.o__Rhizobiales.f__Bradyrhizobiaceae.g__Tardiphaga | 3.83 |
| 15 | k__Bacteria.p__Aquificae.c__Aquificae.o__Aquificales.f__Aquificaceae.g__Hydrogenobaculum | 3.69 |
| 16 | k__Bacteria.p__Proteobacteria.c__Alphaproteobacteria.o__Rhizobiales.f__Rhizobiaceae.g__Rhizobium | 3.69 |
| 17 | k__Bacteria.p__Actinobacteria.c__Actinobacteria.o__Micromonosporales.f__Micromonosporaceae.g__Micromonospora | 3.50 |
| 18 | k__Bacteria.p__Bacteroidetes.c__Cytophagia.o__Cytophagales.f__Flammeovirgaceae.g__Flammeovirga | 3.44 |
| 19 | k__Bacteria.p__Proteobacteria.c__Gammaproteobacteria.o__Enterobacteriales.f__Enterobacteriaceae.g__Proteus | 3.26 |
| 20 | k__Bacteria.p__Proteobacteria.c__Gammaproteobacteria.o__Legionellales.f__Legionellaceae.g__Legionella | 3.25 |

| **Breast Invasive Carcinoma** | | |
| --- | --- | --- |
| **Ranking** | **Microbial Types** | **Contribution (×10^­-3^)** |
| 1 | k__Bacteria.p__Proteobacteria.c__Gammaproteobacteria.o__Aeromonadales.f__Succinivibrionaceae.g__Succinimonas | 5.44 |
| 2 | k__Bacteria.p__Proteobacteria.c__Alphaproteobacteria.o__Rhodospirillales.f__Acetobacteraceae.g__Roseomonas | 4.80 |
| 3 | k__Bacteria.p__Firmicutes.c__Bacilli.o__Bacillales.g__Acidibacillus | 4.78 |
| 4 | k__Bacteria.p__Verrucomicrobia.c__Opitutae.o__Opitutales.f__Opitutaceae.g__Diplosphaera | 4.10 |
| 5 | k__Bacteria.p__Firmicutes.c__Clostridia.o__Clostridiales.f__Lachnospiraceae.g__Lachnobacterium | 4.06 |
| 6 | k__Bacteria.p__Actinobacteria.c__Coriobacteriia.o__Coriobacteriales.f__Atopobiaceae.g__Atopobium | 3.76 |
| 7 | k__Viruses.f__Hypoviridae.g__Hypovirus | 3.53 |
| 8 | k__Bacteria.p__Proteobacteria.c__Gammaproteobacteria.o__Alteromonadales.f__Pseudoalteromonadaceae.g__Algicola | 3.46 |
| 9 | k__Bacteria.p__Bacteroidetes.c__Bacteroidia.o__Bacteroidales.f__Prolixibacteraceae.g__Sunxiuqinia | 3.36 |
| 10 | k__Viruses.f__Nimaviridae.g__Whispovirus | 3.29 |
| 11 | k__Bacteria.p__Proteobacteria.c__Alphaproteobacteria.o__Pelagibacterales.f__Pelagibacteraceae.g__Candidatus_Pelagibacter | 3.28 |
| 12 | k__Bacteria.p__Bacteroidetes.c__Chitinophagia.o__Chitinophagales.f__Saprospiraceae.g__Lewinella | 3.23 |
| 13 | k__Bacteria.p__Proteobacteria.c__Gammaproteobacteria.o__Oceanospirillales.f__Alcanivoracaceae.g__Alcanivorax | 3.20 |
| 14 | k__Bacteria.p__Bacteroidetes.c__Flavobacteriia.o__Flavobacteriales.f__Flavobacteriaceae.g__Ornithobacterium | 3.19 |
| 15 | k__Bacteria.p__Proteobacteria.c__Alphaproteobacteria.o__Rhizobiales.f__Hyphomicrobiaceae.g__Rhodomicrobium | 3.18 |
| 16 | k__Bacteria.p__Firmicutes.c__Clostridia.o__Clostridiales.f__Peptostreptococcaceae.g__Paraclostridium | 3.17 |
| 17 | k__Bacteria.p__Proteobacteria.c__Gammaproteobacteria.o__Oceanospirillales.f__Oceanospirillaceae.g__Marinomonas | 2.95 |
| 18 | k__Bacteria.p__Firmicutes.c__Bacilli.o__Lactobacillales.f__Lactobacillaceae.g__Lactobacillus | 2.95 |
| 19 | k__Viruses.f__Bunyaviridae.g__Tospovirus | 2.88 |
| 20 | k__Bacteria.p__Proteobacteria.c__Betaproteobacteria.o__Burkholderiales.f__Oxalobacteraceae.g__Collimonas | 2.83 |

| **Cervical Squamous Cell Carcinoma and Endocervical Adenocarcinoma** | | |
| --- | --- | --- |
| **Ranking** | **Microbial Types** | **Contribution (×10^­-3^)** |
| 1 | k__Bacteria.p__Proteobacteria.c__Alphaproteobacteria.o__Rhodobacterales.f__Rhodobacteraceae.g__Roseobacter | 18.33 |
| 2 | k__Bacteria.p__Firmicutes.c__Bacilli.o__Bacillales.f__Planococcaceae.g__Paenisporosarcina | 16.61 |
| 3 | k__Bacteria.p__Actinobacteria.c__Actinobacteria.o__Micrococcales.f__Intrasporangiaceae.g__Terrabacter | 13.85 |
| 4 | k__Bacteria.p__Proteobacteria.c__Betaproteobacteria.o__Burkholderiales.f__Oxalobacteraceae.g__Herbaspirillum | 12.50 |
| 5 | k__Bacteria.p__Proteobacteria.c__Alphaproteobacteria.o__Rickettsiales.f__Anaplasmataceae.g__Ehrlichia | 11.75 |
| 6 | k__Bacteria.p__Firmicutes.c__Clostridia.o__Clostridiales.g__Epulopiscium | 11.65 |
| 7 | k__Viruses.f__Astroviridae.g__Avastrovirus | 10.89 |
| 8 | k__Bacteria.p__Cyanobacteria.o__Prochlorales.f__Prochlorococcaceae.g__Prochlorococcus | 10.87 |
| 9 | k__Bacteria.p__Proteobacteria.c__Gammaproteobacteria.o__Pasteurellales.f__Pasteurellaceae.g__Aggregatibacter | 9.98 |
| 10 | k__Viruses.f__Partitiviridae.g__Alphapartitivirus | 9.51 |
| 11 | k__Bacteria.p__Firmicutes.c__Clostridia.o__Clostridiales.f__Ruminococcaceae.g__Acetivibrio | 9.49 |
| 12 | k__Bacteria.p__Proteobacteria.c__Betaproteobacteria.g__Candidatus_Nasuia | 9.49 |
| 13 | k__Bacteria.p__Bacteroidetes.c__Flavobacteriia.o__Flavobacteriales.f__Flavobacteriaceae.g__Xanthomarina | 9.34 |
| 14 | k__Bacteria.p__Proteobacteria.c__Alphaproteobacteria.o__Rhizobiales.f__Brucellaceae.g__Ochrobactrum | 8.49 |
| 15 | k__Bacteria.p__Proteobacteria.c__Alphaproteobacteria.o__Rickettsiales.f__Anaplasmataceae.g__Anaplasma | 8.32 |
| 16 | k__Bacteria.p__Proteobacteria.c__Betaproteobacteria.o__Burkholderiales.f__Alcaligenaceae.g__Alcaligenes | 7.83 |
| 17 | k__Bacteria.p__Proteobacteria.c__Alphaproteobacteria.o__Rhodospirillales.f__Rhodospirillaceae.g__Fodinicurvata | 7.29 |
| 18 | k__Bacteria.p__Proteobacteria.c__Deltaproteobacteria.o__Desulfobacterales.f__Desulfobacteraceae.g__Desulfatibacillum | 7.26 |
| 19 | k__Bacteria.p__Tenericutes.c__Mollicutes.o__Mycoplasmatales.f__Mycoplasmataceae.g__Mycoplasma | 6.96 |
| 20 | k__Bacteria.p__Proteobacteria.c__Gammaproteobacteria.o__Enterobacteriales.f__Enterobacteriaceae.g__Trabulsiella | 6.90 |

| **Colon Adenocarcinoma** | | |
| --- | --- | --- |
| **Ranking** | **Microbial Types** | **Contribution (×10^­-3^)** |
| 1 | k__Bacteria.p__Proteobacteria.c__Alphaproteobacteria.o__Rhizobiales.f__Methylocystaceae.g__Methylocystis | 3.03 |
| 2 | k__Bacteria.p__Cyanobacteria.o__Nostocales.f__Rivulariaceae.g__Calothrix | 3.01 |
| 3 | k__Archaea.p__Crenarchaeota.c__Thermoprotei.o__Sulfolobales.f__Sulfolobaceae.g__Sulfolobus | 3.00 |
| 4 | k__Bacteria.p__Firmicutes.c__Bacilli.o__Bacillales.g__Acidibacillus | 2.99 |
| 5 | k__Bacteria.p__Firmicutes.c__Clostridia.o__Thermoanaerobacterales.f__Thermoanaerobacterales_Family_III._Incertae_Sedis.g__Caldicellulosiruptor | 2.92 |
| 6 | k__Bacteria.p__Proteobacteria.c__Betaproteobacteria.o__Burkholderiales.f__Alcaligenaceae.g__Brackiella | 2.90 |
| 7 | k__Archaea.p__Euryarchaeota.c__Methanobacteria.o__Methanobacteriales.f__Methanobacteriaceae.g__Methanobacterium | 2.84 |
| 8 | k__Viruses.f__Baculoviridae.g__Alphabaculovirus | 2.80 |
| 9 | k__Viruses.f__Bunyaviridae.g__Tospovirus | 2.79 |
| 10 | k__Bacteria.p__Acidobacteria.c__Holophagae.o__Holophagales.f__Holophagaceae.g__Holophaga | 2.71 |
| 11 | k__Bacteria.p__Bacteroidetes.c__Flavobacteriia.o__Flavobacteriales.f__Flavobacteriaceae.g__Tenacibaculum | 2.70 |
| 12 | k__Archaea.p__Euryarchaeota.c__Methanococci.o__Methanococcales.f__Methanocaldococcaceae.g__Methanocaldococcus | 2.67 |
| 13 | k__Viruses.o__Herpesvirales.f__Herpesviridae.g__Varicellovirus | 2.64 |
| 14 | k__Bacteria.p__Proteobacteria.c__Gammaproteobacteria.o__Pasteurellales.f__Pasteurellaceae.g__Necropsobacter | 2.64 |
| 15 | k__Bacteria.p__Proteobacteria.c__Gammaproteobacteria.o__Enterobacteriales.f__Enterobacteriaceae.g__Cosenzaea | 2.64 |
| 16 | k__Bacteria.p__Bacteroidetes.c__Bacteroidia.o__Bacteroidales.f__Porphyromonadaceae.g__Porphyromonas | 2.64 |
| 17 | k__Archaea.p__Euryarchaeota.c__Halobacteria.o__Natrialbales.f__Natrialbaceae.g__Halovivax | 2.52 |
| 18 | k__Viruses.f__Papillomaviridae.g__Alphapapillomavirus | 2.48 |
| 19 | k__Bacteria.p__Cyanobacteria.o__Chroococcales.g__Crocosphaera | 2.46 |
| 20 | k__Bacteria.p__Proteobacteria.c__Alphaproteobacteria.o__Sphingomonadales.f__Sphingomonadaceae.g__Sphingopyxis | 2.45 |

| **Esophageal Aarcinoma** | | |
| --- | --- | --- |
| **Ranking** | **Microbial Types** | **Contribution (×10^­-3^)** |
| 1 | k__Viruses.f__Phycodnaviridae.g__Prymnesiovirus | 12.99 |
| 2 | k__Bacteria.p__Nitrospirae.c__Nitrospira.o__Nitrospirales.f__Nitrospiraceae.g__Nitrospira | 11.76 |
| 3 | k__Bacteria.p__Firmicutes.c__Bacilli.o__Bacillales.g__Acidibacillus | 10.66 |
| 4 | k__Bacteria.p__Bacteroidetes.c__Chitinophagia.o__Chitinophagales.f__Chitinophagaceae.g__Niastella | 8.19 |
| 5 | k__Bacteria.p__Actinobacteria.c__Actinobacteria.o__Pseudonocardiales.f__Pseudonocardiaceae.g__Alloactinosynnema | 7.52 |
| 6 | k__Bacteria.p__Proteobacteria.c__Deltaproteobacteria.o__Desulfuromonadales.f__Desulfuromonadaceae.g__Pelobacter | 7.07 |
| 7 | k__Bacteria.p__Rhodothermaeota.c__Balneolia.o__Balneolales.f__Balneolaceae.g__Gracilimonas | 6.87 |
| 8 | k__Bacteria.p__Actinobacteria.c__Actinobacteria.o__Micrococcales.f__Microbacteriaceae.g__Pseudoclavibacter | 6.51 |
| 9 | k__Bacteria.p__Thermotogae.c__Thermotogae.o__Thermotogales.f__Fervidobacteriaceae.g__Fervidobacterium | 6.14 |
| 10 | k__Bacteria.p__Proteobacteria.c__Deltaproteobacteria.o__Myxococcales.f__Anaeromyxobacteraceae.g__Anaeromyxobacter | 5.90 |
| 11 | k__Bacteria.p__Proteobacteria.c__Gammaproteobacteria.o__Chromatiales.f__Chromatiaceae.g__Thiorhodococcus | 5.55 |
| 12 | k__Bacteria.p__Elusimicrobia.c__Endomicrobia.o__Endomicrobiales.f__Endomicrobiaceae.g__Endomicrobium | 5.40 |
| 13 | k__Bacteria.p__Firmicutes.c__Erysipelotrichia.o__Erysipelotrichales.f__Erysipelotrichaceae.g__Holdemania | 5.23 |
| 14 | k__Bacteria.p__Proteobacteria.c__Alphaproteobacteria.o__Rhizobiales.f__Phyllobacteriaceae.g__Aminobacter | 4.82 |
| 15 | k__Bacteria.p__Proteobacteria.c__Gammaproteobacteria.o__Enterobacteriales.f__Enterobacteriaceae.g__Obesumbacterium | 4.68 |
| 16 | k__Bacteria.p__Firmicutes.c__Clostridia.o__Clostridiales.f__Ruminococcaceae.g__Pseudobacteroides | 4.64 |
| 17 | k__Bacteria.p__Actinobacteria.c__Actinobacteria.o__Micrococcales.f__Microbacteriaceae.g__Agromyces | 4.61 |
| 18 | k__Viruses.f__Nimaviridae.g__Whispovirus | 4.20 |
| 19 | k__Bacteria.p__Proteobacteria.c__Gammaproteobacteria.o__Chromatiales.f__Chromatiaceae.g__Marichromatium | 4.05 |
| 20 | k__Bacteria.p__Proteobacteria.c__Gammaproteobacteria.o__Legionellales.f__Coxiellaceae.g__Diplorickettsia | 4.03 |

| **Head and Neck Squamous Cell Carcinoma** | | |
| --- | --- | --- |
| **Ranking** | **Microbial Types** | **Contribution (×10^­-3^)** |
| 1 | k__Viruses.f__Papillomaviridae.g__Alphapapillomavirus | 14.64 |
| 2 | k__Bacteria.p__Firmicutes.c__Clostridia.o__Clostridiales.f__Clostridiaceae.g__Clostridium | 11.88 |
| 3 | k__Bacteria.p__Firmicutes.c__Bacilli.o__Bacillales.g__Acidibacillus | 7.10 |
| 4 | k__Bacteria.p__Proteobacteria.c__Alphaproteobacteria.o__Rhizobiales.f__Hyphomicrobiaceae.g__Rhodomicrobium | 5.55 |
| 5 | k__Bacteria.p__Cyanobacteria.o__Chroococcales.g__Synechococcus | 5.48 |
| 6 | k__Archaea.p__Euryarchaeota.c__Halobacteria.o__Natrialbales.f__Natrialbaceae.g__Natrialba | 5.43 |
| 7 | k__Bacteria.p__Bacteroidetes.c__Cytophagia.o__Cytophagales.f__Flammeovirgaceae.g__Flammeovirga | 5.17 |
| 8 | k__Archaea.p__Thaumarchaeota.g__Candidatus_Nitrosopelagicus | 4.97 |
| 9 | k__Bacteria.p__Bacteroidetes.c__Bacteroidia.o__Bacteroidales.f__Prevotellaceae.g__Hallella | 4.91 |
| 10 | k__Viruses.f__Polyomaviridae.g__Polyomavirus | 4.68 |
| 11 | k__Bacteria.p__Proteobacteria.c__Alphaproteobacteria.o__Rhizobiales.f__Methylobacteriaceae.g__Microvirga | 4.48 |
| 12 | k__Archaea.p__Euryarchaeota.c__Methanomicrobia.o__Methanocellales.f__Methanocellaceae.g__Methanocella | 4.44 |
| 13 | k__Bacteria.p__Proteobacteria.c__Epsilonproteobacteria.o__Campylobacterales.f__Campylobacteraceae.g__Campylobacter | 4.33 |
| 14 | k__Bacteria.p__Firmicutes.c__Tissierellia.o__Tissierellales.f__Peptoniphilaceae.g__Parvimonas | 4.15 |
| 15 | k__Bacteria.p__Proteobacteria.c__Deltaproteobacteria.o__Myxococcales.f__Myxococcaceae.g__Myxococcus | 3.94 |
| 16 | k__Bacteria.p__Bacteroidetes.c__Flavobacteriia.o__Flavobacteriales.f__Flavobacteriaceae.g__Lacinutrix | 3.86 |
| 17 | k__Bacteria.p__Proteobacteria.c__Gammaproteobacteria.o__Alteromonadales.f__Colwelliaceae.g__Colwellia | 3.83 |
| 18 | k__Bacteria.p__Cyanobacteria.o__Chroococcales.g__Chamaesiphon | 3.63 |
| 19 | k__Bacteria.p__Spirochaetes.c__Spirochaetia.o__Spirochaetales.f__Spirochaetaceae.g__Treponema | 3.61 |
| 20 | k__Bacteria.p__Bacteroidetes.c__Flavobacteriia.o__Flavobacteriales.f__Flavobacteriaceae.g__Flavobacterium | 3.58 |

| **Kidney Chromophobe** | | |
| --- | --- | --- |
| **Ranking** | **Microbial Types** | **Contribution (×10^­-3^)** |
| 1 | k__Bacteria.p__Firmicutes.c__Bacilli.o__Bacillales.f__Bacillaceae.g__Hydrogenibacillus | 21.04 |
| 2 | k__Bacteria.p__Firmicutes.c__Clostridia.o__Clostridiales.g__Flavonifractor | 19.65 |
| 3 | k__Archaea.p__Crenarchaeota.c__Thermoprotei.o__Sulfolobales.f__Sulfolobaceae.g__Sulfolobus | 18.72 |
| 4 | k__Bacteria.p__Bacteroidetes.c__Flavobacteriia.o__Flavobacteriales.f__Flavobacteriaceae.g__Psychroserpens | 18.67 |
| 5 | k__Bacteria.p__Tenericutes.c__Mollicutes.o__Mycoplasmatales.f__Mycoplasmataceae.g__Mycoplasma | 15.52 |
| 6 | k__Bacteria.p__Proteobacteria.c__Alphaproteobacteria.o__Rhizobiales.f__Brucellaceae.g__Brucella | 15.49 |
| 7 | k__Bacteria.p__Firmicutes.c__Bacilli.o__Bacillales.f__Listeriaceae.g__Listeria | 14.97 |
| 8 | k__Bacteria.p__Proteobacteria.c__Alphaproteobacteria.o__Rhizobiales.f__Methylobacteriaceae.g__Microvirga | 13.83 |
| 9 | k__Bacteria.p__Bacteroidetes.c__Flavobacteriia.o__Flavobacteriales.f__Flavobacteriaceae.g__Apibacter | 13.63 |
| 10 | k__Bacteria.p__Proteobacteria.c__Betaproteobacteria.o__Neisseriales.f__Neisseriaceae.g__Neisseria | 13.58 |
| 11 | k__Archaea.p__Euryarchaeota.c__Methanobacteria.o__Methanobacteriales.f__Methanobacteriaceae.g__Methanobrevibacter | 13.45 |
| 12 | k__Bacteria.p__Proteobacteria.c__Alphaproteobacteria.o__Caulobacterales.f__Caulobacteraceae.g__Caulobacter | 12.87 |
| 13 | k__Bacteria.p__Chlamydiae.c__Chlamydiia.o__Chlamydiales.f__Waddliaceae.g__Waddlia | 12.23 |
| 14 | k__Viruses.f__Bunyaviridae.g__Tospovirus | 11.53 |
| 15 | k__Bacteria.p__Spirochaetes.c__Spirochaetia.o__Spirochaetales.f__Spirochaetaceae.g__Treponema | 11.27 |
| 16 | k__Bacteria.p__Firmicutes.c__Clostridia.o__Thermoanaerobacterales.f__Thermoanaerobacteraceae.g__Thermoanaerobacter | 11.03 |
| 17 | k__Bacteria.p__Spirochaetes.c__Spirochaetia.o__Spirochaetales.f__Borreliaceae.g__Borrelia | 10.78 |
| 18 | k__Bacteria.p__Cyanobacteria.o__Chroococcales.g__Cyanothece | 10.35 |
| 19 | k__Bacteria.p__Proteobacteria.c__Gammaproteobacteria.o__Enterobacteriales.f__Enterobacteriaceae.g__Yersinia | 8.90 |
| 20 | k__Bacteria.p__Proteobacteria.c__Gammaproteobacteria.o__Legionellales.f__Legionellaceae.g__Legionella | 8.81 |

| **Kidney Renal Clear Cell Carcinoma** | | |
| --- | --- | --- |
| **Ranking** | **Microbial Types** | **Contribution (×10^­-3^)** |
| 1 | k__Archaea.p__Thaumarchaeota.g__Candidatus_Nitrosopelagicus | 21.17 |
| 2 | k__Bacteria.p__Fibrobacteres.c__Chitinivibrionia.o__Chitinivibrionales.f__Chitinivibrionaceae.g__Chitinivibrio | 11.84 |
| 3 | k__Bacteria.p__Proteobacteria.c__Deltaproteobacteria.o__Desulfobacterales.f__Desulfobulbaceae.g__Desulfotalea | 10.03 |
| 4 | k__Bacteria.p__Proteobacteria.c__Alphaproteobacteria.o__Rhodobacterales.f__Hyphomonadaceae.g__Maricaulis | 8.80 |
| 5 | k__Viruses.o__Herpesvirales.f__Herpesviridae.g__Simplexvirus | 7.96 |
| 6 | k__Bacteria.p__Proteobacteria.c__Gammaproteobacteria.o__Aeromonadales.f__Aeromonadaceae.g__Aeromonas | 7.02 |
| 7 | k__Bacteria.p__Firmicutes.c__Bacilli.o__Lactobacillales.f__Streptococcaceae.g__Streptococcus | 6.66 |
| 8 | k__Bacteria.p__Proteobacteria.c__Gammaproteobacteria.o__Chromatiales.f__Chromatiaceae.g__Rheinheimera | 6.01 |
| 9 | k__Bacteria.p__Firmicutes.c__Bacilli.o__Lactobacillales.f__Carnobacteriaceae.g__Carnobacterium | 5.83 |
| 10 | k__Bacteria.p__Bacteroidetes.c__Cytophagia.o__Cytophagales.f__Cyclobacteriaceae.g__Indibacter | 5.73 |
| 11 | k__Bacteria.p__Actinobacteria.c__Actinobacteria.o__Micrococcales.f__Brevibacteriaceae.g__Brevibacterium | 5.70 |
| 12 | k__Bacteria.p__Proteobacteria.c__Gammaproteobacteria.o__Vibrionales.f__Vibrionaceae.g__Vibrio | 5.17 |
| 13 | k__Bacteria.p__Proteobacteria.c__Deltaproteobacteria.o__Myxococcales.f__Polyangiaceae.g__Sorangium | 5.15 |
| 14 | k__Bacteria.p__Actinobacteria.c__Coriobacteriia.o__Coriobacteriales.f__Atopobiaceae.g__Olsenella | 4.84 |
| 15 | k__Bacteria.p__Cyanobacteria.o__Stigonematales.g__Mastigocoleus | 4.72 |
| 16 | k__Bacteria.p__Proteobacteria.c__Betaproteobacteria.o__Burkholderiales.f__Comamonadaceae.g__Pelomonas | 4.61 |
| 17 | k__Viruses.f__Astroviridae.g__Avastrovirus | 4.52 |
| 18 | k__Bacteria.p__Bacteroidetes.c__Flavobacteriia.o__Flavobacteriales.f__Flavobacteriaceae.g__Apibacter | 4.34 |
| 19 | k__Bacteria.p__Firmicutes.c__Clostridia.o__Clostridiales.f__Lachnospiraceae.g__Lachnoclostridium | 4.27 |
| 20 | k__Viruses.f__Retroviridae.g__Gammaretrovirus | 4.12 |

| **Kidney Renal Papillary Cell Carcinoma** | | |
| --- | --- | --- |
| **Ranking** | **Microbial Types** | **Contribution (×10^­-3^)** |
| 1 | k__Bacteria.p__Proteobacteria.c__Gammaproteobacteria.o__Alteromonadales.f__Pseudoalteromonadaceae.g__Algicola | 11.7108011 |
| 2 | k__Viruses.o__Herpesvirales.f__Alloherpesviridae.g__Cyprinivirus | 8.62431473 |
| 3 | k__Bacteria.p__Proteobacteria.c__Alphaproteobacteria.o__Rhodobacterales.f__Rhodobacteraceae.g__Labrenzia | 8.34038177 |
| 4 | k__Bacteria.p__Actinobacteria.c__Actinobacteria.o__Micromonosporales.f__Micromonosporaceae.g__Actinoplanes | 7.784238 |
| 5 | k__Archaea.p__Euryarchaeota.c__Methanomicrobia.o__Methanocellales.f__Methanocellaceae.g__Methanocella | 7.71366602 |
| 6 | k__Bacteria.p__Firmicutes.c__Bacilli.o__Lactobacillales.f__Streptococcaceae.g__Streptococcus | 7.646919 |
| 7 | k__Bacteria.p__Actinobacteria.c__Actinobacteria.o__Corynebacteriales.f__Dietziaceae.g__Dietzia | 7.64410307 |
| 8 | k__Bacteria.p__Proteobacteria.c__Betaproteobacteria.o__Burkholderiales.f__Oxalobacteraceae.g__Herbaspirillum | 7.43062631 |
| 9 | k__Bacteria.p__Cyanobacteria.o__Chroococcales.g__Microcystis | 7.41249171 |
| 10 | k__Bacteria.p__Proteobacteria.c__Gammaproteobacteria.o__Oceanospirillales.f__Halomonadaceae.g__Candidatus_Portiera | 6.84234879 |
| 11 | k__Archaea.p__Euryarchaeota.c__Methanomicrobia.o__Methanosarcinales.f__Methanosarcinaceae.g__Methanomethylovorans | 6.67018242 |
| 12 | k__Viruses.f__Polyomaviridae.g__Polyomavirus | 6.21539773 |
| 13 | k__Bacteria.p__Proteobacteria.c__Alphaproteobacteria.o__Rhodobacterales.f__Rhodobacteraceae.g__Dinoroseobacter | 5.93900701 |
| 14 | k__Bacteria.p__Proteobacteria.c__Alphaproteobacteria.o__Rhodobacterales.f__Rhodobacteraceae.g__Ruegeria | 5.92514555 |
| 15 | k__Viruses.f__Bromoviridae.g__Bromovirus | 5.82568626 |
| 16 | k__Bacteria.p__Proteobacteria.c__Gammaproteobacteria.o__Chromatiales.f__Ectothiorhodospiraceae.g__Arhodomonas | 5.66020941 |
| 17 | k__Bacteria.p__Proteobacteria.c__Betaproteobacteria.o__Burkholderiales.f__Burkholderiaceae.g__Cupriavidus | 5.59681437 |
| 18 | k__Bacteria.p__Actinobacteria.c__Coriobacteriia.o__Coriobacteriales.f__Atopobiaceae.g__Atopobium | 5.50519456 |
| 19 | k__Bacteria.p__Firmicutes.c__Tissierellia.o__Tissierellales.f__Peptoniphilaceae.g__Helcococcus | 5.4626492 |
| 20 | k__Bacteria.p__Proteobacteria.c__Gammaproteobacteria.o__Enterobacteriales.f__Enterobacteriaceae.g__Klebsiella | 5.39175318 |

| **Brain Lower Grade Glioma** | | |
| --- | --- | --- |
| **Ranking** | **Microbial Types** | **Contribution (×10^­-3^)** |
| 1 | k__Bacteria.p__Proteobacteria.c__Betaproteobacteria.o__Burkholderiales.f__Sutterellaceae.g__Sutterella | 23.63 |
| 2 | k__Bacteria.p__Fibrobacteres.c__Chitinivibrionia.o__Chitinivibrionales.f__Chitinivibrionaceae.g__Chitinivibrio | 10.87 |
| 3 | k__Bacteria.p__Bacteroidetes.c__Cytophagia.o__Cytophagales.f__Flammeovirgaceae.g__Flammeovirga | 9.83 |
| 4 | k__Bacteria.p__Firmicutes.c__Clostridia.o__Clostridiales.f__Lachnospiraceae.g__Lachnoclostridium | 9.51 |
| 5 | k__Viruses.o__Herpesvirales.f__Herpesviridae.g__Mardivirus | 9.38 |
| 6 | k__Bacteria.p__Proteobacteria.c__Alphaproteobacteria.o__Rickettsiales.f__Anaplasmataceae.g__Anaplasma | 7.87 |
| 7 | k__Bacteria.p__Proteobacteria.c__Gammaproteobacteria.o__Enterobacteriales.f__Enterobacteriaceae.g__Proteus | 7.49 |
| 8 | k__Bacteria.p__Proteobacteria.c__Alphaproteobacteria.o__Rhodobacterales.f__Rhodobacteraceae.g__Ruegeria | 7.45 |
| 9 | k__Bacteria.p__Chlamydiae.c__Chlamydiia.o__Chlamydiales.f__Chlamydiaceae.g__Chlamydia | 6.59 |
| 10 | k__Bacteria.p__Cyanobacteria.o__Oscillatoriales.g__Kamptonema | 6.06 |
| 11 | k__Bacteria.p__Firmicutes.c__Bacilli.o__Lactobacillales.f__Carnobacteriaceae.g__Carnobacterium | 5.70 |
| 12 | k__Bacteria.p__Actinobacteria.c__Actinobacteria.o__Corynebacteriales.f__Mycobacteriaceae.g__Mycobacterium | 5.60 |
| 13 | k__Bacteria.p__Bacteroidetes.c__Bacteroidia.o__Bacteroidales.f__Bacteroidaceae.g__Bacteroides | 5.50 |
| 14 | k__Bacteria.p__Bacteroidetes.c__Flavobacteriia.o__Flavobacteriales.f__Flavobacteriaceae.g__Apibacter | 5.40 |
| 15 | k__Bacteria.p__Bacteroidetes.c__Flavobacteriia.o__Flavobacteriales.f__Flavobacteriaceae.g__Ornithobacterium | 5.05 |
| 16 | k__Bacteria.p__Proteobacteria.c__Gammaproteobacteria.o__Xanthomonadales.f__Rhodanobacteraceae.g__Luteibacter | 5.03 |
| 17 | k__Bacteria.p__Proteobacteria.c__Betaproteobacteria.o__Neisseriales.f__Chromobacteriaceae.g__Gulbenkiania | 4.57 |
| 18 | k__Viruses.o__Herpesvirales.f__Herpesviridae.g__Proboscivirus | 4.41 |
| 19 | k__Bacteria.p__Bacteroidetes.c__Flavobacteriia.o__Flavobacteriales.f__Flavobacteriaceae.g__Riemerella | 4.30 |
| 20 | k__Bacteria.p__Actinobacteria.c__Actinobacteria.o__Pseudonocardiales.f__Pseudonocardiaceae.g__Alloactinosynnema | 4.09 |

| **Liver Hepatocellular Carcinoma** | | |
| --- | --- | --- |
| **Ranking** | **Microbial Types** | **Contribution (×10^­-3^)** |
| 1 | k__Bacteria.p__Tenericutes.c__Mollicutes.o__Mycoplasmatales.f__Mycoplasmataceae.g__Mycoplasma | 25.07 |
| 2 | k__Bacteria.p__Spirochaetes.c__Spirochaetia.f__Leptospiraceae.g__Leptospira | 22.38 |
| 3 | k__Bacteria.p__Actinobacteria.c__Actinobacteria.o__Micrococcales.f__Microbacteriaceae.g__Curtobacterium | 18.40 |
| 4 | k__Bacteria.p__Proteobacteria.c__Gammaproteobacteria.o__Vibrionales.f__Vibrionaceae.g__Grimontia | 12.99 |
| 5 | k__Bacteria.p__Proteobacteria.c__Acidithiobacillia.o__Acidithiobacillales.f__Acidithiobacillaceae.g__Acidithiobacillus | 12.79 |
| 6 | k__Bacteria.p__Proteobacteria.c__Deltaproteobacteria.o__Desulfobacterales.f__Desulfobacteraceae.g__Desulfatibacillum | 11.31 |
| 7 | k__Bacteria.p__Proteobacteria.c__Betaproteobacteria.o__Burkholderiales.f__Alcaligenaceae.g__Brackiella | 10.04 |
| 8 | k__Bacteria.p__Proteobacteria.c__Gammaproteobacteria.o__Enterobacteriales.f__Enterobacteriaceae.g__Trabulsiella | 9.42 |
| 9 | k__Bacteria.p__Proteobacteria.c__Gammaproteobacteria.o__Pasteurellales.f__Pasteurellaceae.g__Haemophilus | 9.12 |
| 10 | k__Bacteria.p__Actinobacteria.c__Actinobacteria.o__Propionibacteriales.f__Nocardioidaceae.g__Nocardioides | 8.95 |
| 11 | k__Bacteria.p__Bacteroidetes.c__Flavobacteriia.o__Flavobacteriales.f__Flavobacteriaceae.g__Elizabethkingia | 8.73 |
| 12 | k__Archaea.p__Euryarchaeota.c__Methanobacteria.o__Methanobacteriales.f__Methanobacteriaceae.g__Methanobrevibacter | 8.62 |
| 13 | k__Bacteria.p__Actinobacteria.c__Actinobacteria.o__Bifidobacteriales.f__Bifidobacteriaceae.g__Bifidobacterium | 8.03 |
| 14 | k__Bacteria.p__Firmicutes.c__Bacilli.o__Bacillales.f__Bacillaceae.g__Fictibacillus | 8.01 |
| 15 | k__Archaea.p__Euryarchaeota.c__Methanococci.o__Methanococcales.f__Methanococcaceae.g__Methanococcus | 7.42 |
| 16 | k__Bacteria.p__Proteobacteria.c__Alphaproteobacteria.o__Rhodospirillales.f__Acetobacteraceae.g__Acetobacter | 7.15 |
| 17 | k__Bacteria.p__Firmicutes.c__Clostridia.o__Clostridiales.f__Peptostreptococcaceae.g__Clostridioides | 6.76 |
| 18 | k__Bacteria.p__Firmicutes.c__Clostridia.o__Clostridiales.g__Epulopiscium | 6.76 |
| 19 | k__Bacteria.p__Firmicutes.c__Clostridia.o__Thermoanaerobacterales.f__Thermoanaerobacterales_Family_III._Incertae_Sedis.g__Thermoanaerobacterium | 6.58 |
| 20 | k__Bacteria.p__Fusobacteria.c__Fusobacteriia.o__Fusobacteriales.f__Leptotrichiaceae.g__Leptotrichia | 6.43 |

| **Lung Squamous Cell Carcinoma** | | |
| --- | --- | --- |
| **Ranking** | **Microbial Types** | **Contribution (×10^­-3^)** |
| 1 | k__Bacteria.p__Tenericutes.c__Mollicutes.o__Mycoplasmatales.f__Mycoplasmataceae.g__Mycoplasma | 25.07 |
| 2 | k__Bacteria.p__Spirochaetes.c__Spirochaetia.f__Leptospiraceae.g__Leptospira | 22.38 |
| 3 | k__Bacteria.p__Actinobacteria.c__Actinobacteria.o__Micrococcales.f__Microbacteriaceae.g__Curtobacterium | 18.40 |
| 4 | k__Bacteria.p__Proteobacteria.c__Gammaproteobacteria.o__Vibrionales.f__Vibrionaceae.g__Grimontia | 12.99 |
| 5 | k__Bacteria.p__Proteobacteria.c__Acidithiobacillia.o__Acidithiobacillales.f__Acidithiobacillaceae.g__Acidithiobacillus | 12.79 |
| 6 | k__Bacteria.p__Proteobacteria.c__Deltaproteobacteria.o__Desulfobacterales.f__Desulfobacteraceae.g__Desulfatibacillum | 11.31 |
| 7 | k__Bacteria.p__Proteobacteria.c__Betaproteobacteria.o__Burkholderiales.f__Alcaligenaceae.g__Brackiella | 10.04 |
| 8 | k__Bacteria.p__Proteobacteria.c__Gammaproteobacteria.o__Enterobacteriales.f__Enterobacteriaceae.g__Trabulsiella | 9.42 |
| 9 | k__Bacteria.p__Proteobacteria.c__Gammaproteobacteria.o__Pasteurellales.f__Pasteurellaceae.g__Haemophilus | 9.12 |
| 10 | k__Bacteria.p__Actinobacteria.c__Actinobacteria.o__Propionibacteriales.f__Nocardioidaceae.g__Nocardioides | 8.95 |
| 11 | k__Bacteria.p__Bacteroidetes.c__Flavobacteriia.o__Flavobacteriales.f__Flavobacteriaceae.g__Elizabethkingia | 8.73 |
| 12 | k__Archaea.p__Euryarchaeota.c__Methanobacteria.o__Methanobacteriales.f__Methanobacteriaceae.g__Methanobrevibacter | 8.62 |
| 13 | k__Bacteria.p__Actinobacteria.c__Actinobacteria.o__Bifidobacteriales.f__Bifidobacteriaceae.g__Bifidobacterium | 8.03 |
| 14 | k__Bacteria.p__Firmicutes.c__Bacilli.o__Bacillales.f__Bacillaceae.g__Fictibacillus | 8.01 |
| 15 | k__Archaea.p__Euryarchaeota.c__Methanococci.o__Methanococcales.f__Methanococcaceae.g__Methanococcus | 7.42 |
| 16 | k__Bacteria.p__Proteobacteria.c__Alphaproteobacteria.o__Rhodospirillales.f__Acetobacteraceae.g__Acetobacter | 7.15 |
| 17 | k__Bacteria.p__Firmicutes.c__Clostridia.o__Clostridiales.f__Peptostreptococcaceae.g__Clostridioides | 6.76 |
| 18 | k__Bacteria.p__Firmicutes.c__Clostridia.o__Clostridiales.g__Epulopiscium | 6.76 |
| 19 | k__Bacteria.p__Firmicutes.c__Clostridia.o__Thermoanaerobacterales.f__Thermoanaerobacterales_Family_III._Incertae_Sedis.g__Thermoanaerobacterium | 6.58 |
| 20 | k__Bacteria.p__Fusobacteria.c__Fusobacteriia.o__Fusobacteriales.f__Leptotrichiaceae.g__Leptotrichia | 6.43 |

| **Pancreatic Adenocarcinoma** | | |
| --- | --- | --- |
| **Ranking** | **Microbial Types** | **Contribution (×10^­-3^)** |
| 1 | k__Bacteria.p__Proteobacteria.c__Betaproteobacteria.o__Burkholderiales.f__Comamonadaceae.g__Hylemonella | 14.98 |
| 2 | k__Bacteria.p__Bacteroidetes.c__Flavobacteriia.o__Flavobacteriales.f__Flavobacteriaceae.g__Ornithobacterium | 9.06 |
| 3 | k__Bacteria.p__Proteobacteria.c__Gammaproteobacteria.o__Pseudomonadales.f__Moraxellaceae.g__Moraxella | 7.76 |
| 4 | k__Bacteria.p__Proteobacteria.c__Gammaproteobacteria.f__Competibacteraceae.g__Candidatus_Contendobacter | 6.52 |
| 5 | k__Bacteria.p__Proteobacteria.c__Alphaproteobacteria.o__Rhodobacterales.f__Hyphomonadaceae.g__Hyphomonas | 6.22 |
| 6 | k__Bacteria.p__Firmicutes.c__Bacilli.o__Lactobacillales.f__Lactobacillaceae.g__Pediococcus | 6.11 |
| 7 | k__Bacteria.p__Firmicutes.c__Bacilli.o__Bacillales.f__Bacillaceae.g__Hydrogenibacillus | 6.06 |
| 8 | k__Bacteria.p__Proteobacteria.c__Alphaproteobacteria.o__Rhodobacterales.f__Rhodobacteraceae.g__Tropicibacter | 5.76 |
| 9 | k__Bacteria.p__Proteobacteria.c__Betaproteobacteria.o__Burkholderiales.f__Comamonadaceae.g__Alicycliphilus | 5.65 |
| 10 | k__Bacteria.p__Proteobacteria.c__Betaproteobacteria.o__Hydrogenophilales.f__Hydrogenophilaceae.g__Tepidiphilus | 5.56 |
| 11 | k__Bacteria.p__Proteobacteria.c__Betaproteobacteria.o__Burkholderiales.g__Sphaerotilus | 5.43 |
| 12 | k__Bacteria.p__Planctomycetes.c__Planctomycetia.o__Planctomycetales.f__Planctomycetaceae.g__Planctomyces | 5.33 |
| 13 | k__Bacteria.p__Cyanobacteria.o__Nostocales.f__Microchaetaceae.g__Microchaete | 5.20 |
| 14 | k__Bacteria.p__Proteobacteria.c__Gammaproteobacteria.o__Vibrionales.f__Vibrionaceae.g__Photobacterium | 5.16 |
| 15 | k__Viruses.f__Poxviridae.g__Molluscipoxvirus | 5.13 |
| 16 | k__Viruses.f__Retroviridae.g__Gammaretrovirus | 5.07 |
| 17 | k__Bacteria.p__Bacteroidetes.c__Flavobacteriia.o__Flavobacteriales.f__Blattabacteriaceae.g__Blattabacterium | 5.04 |
| 18 | k__Bacteria.p__Proteobacteria.c__Betaproteobacteria.o__Burkholderiales.g__Paucibacter | 4.83 |
| 19 | k__Bacteria.p__Cyanobacteria.o__Nostocales.f__Rivulariaceae.g__Calothrix | 4.69 |
| 20 | k__Bacteria.p__Deinococcus.Thermus.c__Deinococci.o__Thermales.f__Thermaceae.g__Meiothermus | 4.62 |

| **Rectum Adenocarcinoma** | | |
| --- | --- | --- |
| **Ranking** | **Microbial Types** | **Contribution (×10^­-3^)** |
| 1 | k__Bacteria.p__Actinobacteria.c__Actinobacteria.o__Corynebacteriales.f__Dietziaceae.g__Dietzia | 15.57 |
| 2 | k__Viruses.f__Retroviridae.g__Alpharetrovirus | 11.00 |
| 3 | k__Viruses.o__Herpesvirales.f__Herpesviridae.g__Cytomegalovirus | 8.41 |
| 4 | k__Bacteria.p__Actinobacteria.c__Coriobacteriia.o__Coriobacteriales.f__Atopobiaceae.g__Atopobium | 6.39 |
| 5 | k__Bacteria.p__Firmicutes.c__Bacilli.o__Bacillales.f__Planococcaceae.g__Planococcus | 6.24 |
| 6 | k__Bacteria.p__Actinobacteria.c__Actinobacteria.o__Micrococcales.f__Promicromonosporaceae.g__Cellulosimicrobium | 6.15 |
| 7 | k__Bacteria.p__Proteobacteria.c__Gammaproteobacteria.o__Chromatiales.f__Ectothiorhodospiraceae.g__Thioalkalivibrio | 6.01 |
| 8 | k__Bacteria.p__Proteobacteria.c__Deltaproteobacteria.o__Desulfovibrionales.f__Desulfonatronaceae.g__Desulfonatronum | 5.40 |
| 9 | k__Bacteria.p__Proteobacteria.c__Gammaproteobacteria.o__Enterobacteriales.f__Enterobacteriaceae.g__Proteus | 5.11 |
| 10 | k__Bacteria.p__Proteobacteria.c__Betaproteobacteria.o__Burkholderiales.g__Tepidimonas | 5.05 |
| 11 | k__Bacteria.p__Bacteroidetes.c__Cytophagia.o__Cytophagales.f__Cyclobacteriaceae.g__Indibacter | 5.03 |
| 12 | k__Bacteria.p__Proteobacteria.c__Gammaproteobacteria.o__Enterobacteriales.f__Enterobacteriaceae.g__Wigglesworthia | 4.92 |
| 13 | k__Bacteria.p__Proteobacteria.c__Betaproteobacteria.o__Burkholderiales.g__Rhizobacter | 4.89 |
| 14 | k__Viruses.f__Phycodnaviridae.g__Prymnesiovirus | 4.77 |
| 15 | k__Archaea.p__Euryarchaeota.c__Halobacteria.o__Natrialbales.f__Natrialbaceae.g__Natrialba | 4.76 |
| 16 | k__Bacteria.p__Proteobacteria.c__Gammaproteobacteria.o__Xanthomonadales.f__Rhodanobacteraceae.g__Luteibacter | 4.47 |
| 17 | k__Bacteria.p__Proteobacteria.c__Alphaproteobacteria.o__Rhodospirillales.f__Rhodospirillaceae.g__Rhodovibrio | 4.46 |
| 18 | k__Bacteria.p__Actinobacteria.c__Actinobacteria.o__Micrococcales.f__Dermatophilaceae.g__Dermatophilus | 4.42 |
| 19 | k__Bacteria.p__Chloroflexi.c__Anaerolineae.o__Anaerolineales.f__Anaerolineaceae.g__Ornatilinea | 4.41 |
| 20 | k__Bacteria.p__Actinobacteria.c__Actinobacteria.o__Propionibacteriales.f__Nocardioidaceae.g__Nocardioides | 4.32 |

| **Sarcoma** | | |
| --- | --- | --- |
| **Ranking** | **Microbial Types** | **Contribution (×10^­-3^)** |
| 1 | k__Bacteria.p__Actinobacteria.c__Actinobacteria.o__Streptosporangiales.f__Streptosporangiaceae.g__Streptosporangium | 10.82 |
| 2 | k__Bacteria.p__Bacteroidetes.c__Cytophagia.o__Cytophagales.f__Flammeovirgaceae.g__Flammeovirga | 6.37 |
| 3 | k__Bacteria.p__Proteobacteria.c__Gammaproteobacteria.o__Oceanospirillales.f__Oceanospirillaceae.g__Marinomonas | 5.87 |
| 4 | k__Bacteria.p__Proteobacteria.c__Betaproteobacteria.o__Burkholderiales.f__Comamonadaceae.g__Curvibacter | 5.78 |
| 5 | k__Bacteria.p__Firmicutes.c__Clostridia.o__Clostridiales.f__Lachnospiraceae.g__Lachnoclostridium | 5.57 |
| 6 | k__Archaea.p__Thaumarchaeota.g__Candidatus_Nitrosopelagicus | 5.43 |
| 7 | k__Bacteria.p__Actinobacteria.c__Coriobacteriia.o__Coriobacteriales.f__Coriobacteriaceae.g__Collinsella | 4.93 |
| 8 | k__Bacteria.p__Fibrobacteres.c__Chitinivibrionia.o__Chitinivibrionales.f__Chitinivibrionaceae.g__Chitinivibrio | 4.92 |
| 9 | k__Bacteria.p__Firmicutes.c__Bacilli.o__Bacillales.f__Staphylococcaceae.g__Staphylococcus | 4.75 |
| 10 | k__Bacteria.p__Proteobacteria.c__Betaproteobacteria.o__Burkholderiales.f__Comamonadaceae.g__Acidovorax | 4.62 |
| 11 | k__Bacteria.p__Proteobacteria.c__Gammaproteobacteria.o__Enterobacteriales.f__Enterobacteriaceae.g__Proteus | 4.49 |
| 12 | k__Bacteria.p__Actinobacteria.c__Actinobacteria.o__Micrococcales.f__Dermacoccaceae.g__Kytococcus | 4.30 |
| 13 | k__Bacteria.p__Firmicutes.c__Clostridia.o__Clostridiales.g__Flavonifractor | 4.17 |
| 14 | k__Viruses.f__Togaviridae.g__Alphavirus | 4.01 |
| 15 | k__Archaea.p__Euryarchaeota.c__Halobacteria.o__Halobacteriales.f__Halobacteriaceae.g__Halococcus | 3.98 |
| 16 | k__Bacteria.p__Firmicutes.c__Bacilli.o__Bacillales.g__Exiguobacterium | 3.91 |
| 17 | k__Bacteria.p__Proteobacteria.c__Gammaproteobacteria.o__Enterobacteriales.f__Enterobacteriaceae.g__Xenorhabdus | 3.88 |
| 18 | k__Bacteria.p__Actinobacteria.c__Actinobacteria.o__Micrococcales.f__Dermabacteraceae.g__Brachybacterium | 3.88 |
| 19 | k__Bacteria.p__Proteobacteria.c__Gammaproteobacteria.o__Pseudomonadales.f__Moraxellaceae.g__Acinetobacter | 3.86 |
| 20 | k__Bacteria.p__Proteobacteria.c__Alphaproteobacteria.o__Rhizobiales.f__Phyllobacteriaceae.g__Mesorhizobium | 3.73 |

| **Skin Cutaneous Melanoma** | | |
| --- | --- | --- |
| **Ranking** | **Microbial Types** | **Contribution (×10^­-3^)** |
| 1 | k__Bacteria.p__Actinobacteria.c__Actinobacteria.o__Micrococcales.f__Dermabacteraceae.g__Brachybacterium | 21.60 |
| 2 | k__Bacteria.p__Proteobacteria.c__Betaproteobacteria.o__Burkholderiales.f__Comamonadaceae.g__Acidovorax | 20.86 |
| 3 | k__Bacteria.p__Proteobacteria.c__Deltaproteobacteria.o__Desulfobacterales.f__Desulfobacteraceae.g__Desulfococcus | 19.44 |
| 4 | k__Bacteria.p__Firmicutes.c__Bacilli.o__Lactobacillales.f__Lactobacillaceae.g__Pediococcus | 18.36 |
| 5 | k__Bacteria.p__Proteobacteria.c__Betaproteobacteria.o__Burkholderiales.g__Thiomonas | 16.56 |
| 6 | k__Bacteria.p__Proteobacteria.c__Gammaproteobacteria.f__Competibacteraceae.g__Candidatus_Contendobacter | 16.49 |
| 7 | k__Bacteria.p__Bacteroidetes.c__Chitinophagia.o__Chitinophagales.f__Chitinophagaceae.g__Sediminibacterium | 13.29 |
| 8 | k__Bacteria.p__Proteobacteria.c__Betaproteobacteria.o__Neisseriales.f__Chromobacteriaceae.g__Pseudogulbenkiania | 11.42 |
| 9 | k__Bacteria.p__Proteobacteria.c__Alphaproteobacteria.o__Rhizobiales.f__Rhizobiaceae.g__Agrobacterium | 11.22 |
| 10 | k__Bacteria.p__Proteobacteria.c__Betaproteobacteria.o__Burkholderiales.f__Comamonadaceae.g__Pelomonas | 11.08 |
| 11 | k__Bacteria.p__Proteobacteria.c__Gammaproteobacteria.o__Xanthomonadales.f__Xanthomonadaceae.g__Xanthomonas | 11.01 |
| 12 | k__Bacteria.p__Firmicutes.c__Bacilli.o__Bacillales.f__Staphylococcaceae.g__Staphylococcus | 10.45 |
| 13 | k__Bacteria.p__Actinobacteria.c__Actinobacteria.o__Actinomycetales.f__Actinomycetaceae.g__Actinomyces | 10.44 |
| 14 | k__Bacteria.p__Proteobacteria.c__Betaproteobacteria.o__Burkholderiales.f__Comamonadaceae.g__Curvibacter | 9.91 |
| 15 | k__Bacteria.p__Proteobacteria.c__Gammaproteobacteria.o__Enterobacteriales.f__Enterobacteriaceae.g__Enterobacter | 9.72 |
| 16 | k__Bacteria.p__Bacteroidetes.c__Chitinophagia.o__Chitinophagales.f__Saprospiraceae.g__Aureispira | 9.68 |
| 17 | k__Bacteria.p__Proteobacteria.c__Alphaproteobacteria.o__Rhodospirillales.f__Acetobacteraceae.g__Acetobacter | 9.51 |
| 18 | k__Bacteria.p__Proteobacteria.c__Betaproteobacteria.o__Burkholderiales.f__Comamonadaceae.g__Variovorax | 9.27 |
| 19 | k__Bacteria.p__Proteobacteria.c__Betaproteobacteria.o__Neisseriales.f__Chromobacteriaceae.g__Gulbenkiania | 9.22 |
| 20 | k__Bacteria.p__Actinobacteria.c__Actinobacteria.o__Micrococcales.f__Microbacteriaceae.g__Agromyces | 8.77 |

| **Thymoma** | | |
| --- | --- | --- |
| **Ranking** | **Microbial Types** | **Contribution (×10^­-3^)** |
| 1 | k__Viruses.o__Herpesvirales.f__Alloherpesviridae.g__Cyprinivirus | 12.18 |
| 2 | k__Bacteria.p__Bacteroidetes.c__Flavobacteriia.o__Flavobacteriales.f__Flavobacteriaceae.g__Imtechella | 10.19 |
| 3 | k__Bacteria.p__Proteobacteria.c__Gammaproteobacteria.o__Enterobacteriales.f__Enterobacteriaceae.g__Proteus | 9.63 |
| 4 | k__Viruses.f__Phycodnaviridae.g__Phaeovirus | 9.46 |
| 5 | k__Bacteria.p__Bacteroidetes.o__Bacteroidetes_Order_II._Incertae_sedis.f__Rhodothermaceae.g__Rhodothermus | 9.46 |
| 6 | k__Bacteria.p__Firmicutes.c__Clostridia.o__Clostridiales.f__Syntrophomonadaceae.g__Dethiobacter | 8.16 |
| 7 | k__Bacteria.p__Proteobacteria.c__Gammaproteobacteria.o__Enterobacteriales.f__Enterobacteriaceae.g__Phaseolibacter | 8.07 |
| 8 | k__Bacteria.p__Actinobacteria.c__Actinobacteria.o__Micrococcales.f__Microbacteriaceae.g__Frigoribacterium | 7.78 |
| 9 | k__Bacteria.p__Actinobacteria.c__Actinobacteria.o__Micrococcales.f__Promicromonosporaceae.g__Cellulosimicrobium | 7.46 |
| 10 | k__Bacteria.p__Spirochaetes.c__Spirochaetia.o__Brachyspirales.f__Brachyspiraceae.g__Brachyspira | 7.41 |
| 11 | k__Bacteria.p__Bacteroidetes.c__Flavobacteriia.o__Flavobacteriales.f__Flavobacteriaceae.g__Apibacter | 7.03 |
| 12 | k__Bacteria.p__Gemmatimonadetes.c__Gemmatimonadetes.o__Gemmatimonadales.f__Gemmatimonadaceae.g__Gemmatimonas | 6.87 |
| 13 | k__Bacteria.p__Actinobacteria.c__Coriobacteriia.o__Coriobacteriales.f__Atopobiaceae.g__Olsenella | 6.67 |
| 14 | k__Bacteria.p__Firmicutes.c__Bacilli.o__Bacillales.f__Bacillaceae.g__Fictibacillus | 6.60 |
| 15 | k__Bacteria.p__Verrucomicrobia.c__Opitutae.o__Opitutales.f__Opitutaceae.g__Cephaloticoccus | 6.45 |
| 16 | k__Bacteria.p__Cyanobacteria.o__Chroococcales.g__Chamaesiphon | 6.31 |
| 17 | k__Viruses.o__Tymovirales.f__Tymoviridae.g__Tymovirus | 6.18 |
| 18 | k__Bacteria.p__Proteobacteria.c__Gammaproteobacteria.o__Alteromonadales.f__Alteromonadaceae.g__Glaciecola | 6.17 |
| 19 | k__Bacteria.p__Firmicutes.c__Bacilli.o__Lactobacillales.f__Aerococcaceae.g__Aerococcus | 5.77 |
| 20 | k__Bacteria.p__Bacteroidetes.c__Bacteroidia.o__Bacteroidales.f__Bacteroidaceae.g__Bacteroides | 5.72 |

| **Uterine Corpus Endometrial Carcinoma** | | |
| --- | --- | --- |
| **Ranking** | **Microbial Types** | **Contribution (×10^­-3^)** |
| 1 | k__Bacteria.p__Actinobacteria.c__Actinobacteria.o__Micrococcales.f__Microbacteriaceae.g__Plantibacter | 11.45 |
| 2 | k__Bacteria.p__Firmicutes.c__Bacilli.o__Bacillales.g__Acidibacillus | 9.21 |
| 3 | k__Bacteria.p__Proteobacteria.c__Betaproteobacteria.o__Rhodocyclales.f__Rhodocyclaceae.g__Thauera | 7.31 |
| 4 | k__Bacteria.p__Proteobacteria.c__Gammaproteobacteria.o__Chromatiales.f__Chromatiaceae.g__Rheinheimera | 7.02 |
| 5 | k__Bacteria.p__Proteobacteria.c__Alphaproteobacteria.o__Rhizobiales.f__Aurantimonadaceae.g__Aureimonas | 6.14 |
| 6 | k__Bacteria.p__Proteobacteria.c__Gammaproteobacteria.o__Enterobacteriales.f__Enterobacteriaceae.g__Enterobacter | 6.02 |
| 7 | k__Bacteria.p__Proteobacteria.c__Alphaproteobacteria.o__Rhodobacterales.f__Rhodobacteraceae.g__Sulfitobacter | 5.45 |
| 8 | k__Bacteria.p__Firmicutes.c__Clostridia.o__Clostridiales.f__Peptostreptococcaceae.g__Paeniclostridium | 5.38 |
| 9 | k__Bacteria.p__Proteobacteria.c__Alphaproteobacteria.o__Rhodospirillales.f__Acetobacteraceae.g__Saccharibacter | 4.81 |
| 10 | k__Bacteria.p__Proteobacteria.c__Betaproteobacteria.g__Candidatus_Accumulibacter | 4.39 |
| 11 | k__Viruses.f__Nimaviridae.g__Whispovirus | 4.02 |
| 12 | k__Bacteria.p__Proteobacteria.c__Betaproteobacteria.o__Burkholderiales.f__Burkholderiaceae.g__Limnobacter | 3.97 |
| 13 | k__Bacteria.p__Proteobacteria.c__Betaproteobacteria.o__Rhodocyclales.f__Rhodocyclaceae.g__Dechloromonas | 3.91 |
| 14 | k__Bacteria.p__Cyanobacteria.o__Chroococcales.g__Cyanothece | 3.83 |
| 15 | k__Bacteria.p__Proteobacteria.c__Alphaproteobacteria.o__Sphingomonadales.f__Sphingomonadaceae.g__Sphingopyxis | 3.73 |
| 16 | k__Bacteria.p__Proteobacteria.c__Alphaproteobacteria.o__Rhizobiales.f__Bradyrhizobiaceae.g__Rhodopseudomonas | 3.65 |
| 17 | k__Bacteria.p__Proteobacteria.c__Gammaproteobacteria.o__Enterobacteriales.f__Enterobacteriaceae.g__Erwinia | 3.63 |
| 18 | k__Bacteria.p__Proteobacteria.c__Betaproteobacteria.o__Burkholderiales.f__Comamonadaceae.g__Diaphorobacter | 3.62 |
| 19 | k__Bacteria.p__Actinobacteria.c__Actinobacteria.o__Propionibacteriales.f__Nocardioidaceae.g__Nocardioides | 3.54 |
| 20 | k__Bacteria.p__Actinobacteria.c__Actinobacteria.o__Micrococcales.f__Intrasporangiaceae.g__Terrabacter | 3.42 |

| **Uveal Melanoma** | | |
| --- | --- | --- |
| **Ranking** | **Microbial Types** | **Contribution (×10^­-3^)** |
| 1 | k__Bacteria.p__Proteobacteria.c__Alphaproteobacteria.o__Rhodobacterales.f__Rhodobacteraceae.g__Rhodovulum | 16.65 |
| 2 | k__Bacteria.p__Actinobacteria.c__Actinobacteria.o__Micrococcales.f__Micrococcaceae.g__Nesterenkonia | 13.38 |
| 3 | k__Viruses.f__Nimaviridae.g__Whispovirus | 12.19 |
| 4 | k__Bacteria.p__Proteobacteria.c__Betaproteobacteria.o__Rhodocyclales.f__Rhodocyclaceae.g__Dechloromonas | 11.44 |
| 5 | k__Bacteria.p__Proteobacteria.c__Alphaproteobacteria.o__Rhizobiales.f__Rhizobiaceae.g__Ensifer | 8.47 |
| 6 | k__Bacteria.p__Proteobacteria.c__Betaproteobacteria.o__Rhodocyclales.f__Rhodocyclaceae.g__Aromatoleum | 8.15 |
| 7 | k__Bacteria.p__Proteobacteria.c__Alphaproteobacteria.o__Rhizobiales.f__Phyllobacteriaceae.g__Nitratireductor | 7.72 |
| 8 | k__Bacteria.p__Proteobacteria.c__Gammaproteobacteria.o__Alteromonadales.f__Pseudoalteromonadaceae.g__Algicola | 7.54 |
| 9 | k__Bacteria.p__Proteobacteria.c__Gammaproteobacteria.o__Pseudomonadales.f__Pseudomonadaceae.g__Pseudomonas | 6.94 |
| 10 | k__Bacteria.p__Actinobacteria.c__Actinobacteria.o__Micrococcales.f__Dermacoccaceae.g__Kytococcus | 6.82 |
| 11 | k__Bacteria.p__Proteobacteria.c__Alphaproteobacteria.o__Rhodospirillales.f__Rhodospirillaceae.g__Skermanella | 6.65 |
| 12 | k__Bacteria.p__Firmicutes.c__Clostridia.o__Clostridiales.f__Eubacteriaceae.g__Acetobacterium | 6.37 |
| 13 | k__Bacteria.p__Proteobacteria.c__Alphaproteobacteria.o__Rhodobacterales.f__Rhodobacteraceae.g__Roseobacter | 5.95 |
| 14 | k__Bacteria.p__Bacteroidetes.c__Cytophagia.o__Cytophagales.f__Flammeovirgaceae.g__Flammeovirga | 5.93 |
| 15 | k__Bacteria.p__Firmicutes.c__Bacilli.o__Bacillales.f__Paenibacillaceae.g__Cohnella | 5.63 |
| 16 | k__Archaea.p__Euryarchaeota.c__Halobacteria.o__Natrialbales.f__Natrialbaceae.g__Halovivax | 5.37 |
| 17 | k__Bacteria.p__Bacteroidetes.c__Flavobacteriia.o__Flavobacteriales.f__Flavobacteriaceae.g__Maribacter | 5.36 |
| 18 | k__Bacteria.p__Proteobacteria.c__Alphaproteobacteria.o__Rhizobiales.f__Rhizobiaceae.g__Sinorhizobium | 4.98 |
| 19 | k__Bacteria.p__Actinobacteria.c__Actinobacteria.o__Propionibacteriales.f__Propionibacteriaceae.g__Propionibacterium | 4.98 |
| 20 | k__Bacteria.p__Proteobacteria.c__Betaproteobacteria.o__Rhodocyclales.f__Rhodocyclaceae.g__Thauera | 4.90 |

**Supplementary Table 3. Top 20 outstanding host gene with highest contribution of each cancer in subtype prediction.**

| **Adrenocortical Carcinoma** | | |
| --- | --- | --- |
| **Ranking** | **Host Genes** | **Contribution (×10^­-3^)** |
| 1 | LHX4-AS1 | 18.91 |
| 2 | PPRC1 | 18.30 |
| 3 | CRTC2 | 18.07 |
| 4 | 5-Sep | 15.29 |
| 5 | B4GALT2 | 15.21 |
| 6 | CD97 | 12.46 |
| 7 | ZMYND19 | 12.29 |
| 8 | GDI2 | 11.35 |
| 9 | DEDD | 10.13 |
| 10 | SMAD2 | 9.89 |
| 11 | CEP76 | 9.13 |
| 12 | SPNS3 | 8.68 |
| 13 | SF3B4 | 8.57 |
| 14 | SUV39H2 | 8.25 |
| 15 | TRMT2A | 8.14 |
| 16 | HDAC4 | 8.10 |
| 17 | WDR77 | 8.09 |
| 18 | ILKAP | 7.82 |
| 19 | SLC29A4 | 7.63 |
| 20 | FAM177A1 | 7.37 |

| **Bladder Urothelial Carcinoma** | | |
| --- | --- | --- |
| **Ranking** | **Host Genes** | **Contribution (×10^­-3^)** |
| 1 | GNB4 | 11.34 |
| 2 | LOX | 11.22 |
| 3 | TGFB3 | 10.18 |
| 4 | CHST11 | 8.13 |
| 5 | ECM1 | 7.85 |
| 6 | MT2A | 7.06 |
| 7 | MN1 | 6.92 |
| 8 | KATNAL1 | 6.85 |
| 9 | COL8A2 | 6.78 |
| 10 | SWSAP1 | 6.59 |
| 11 | FLNC | 6.53 |
| 12 | TNFAIP8L3 | 6.30 |
| 13 | TCP11L1 | 6.03 |
| 14 | ZFHX4 | 5.64 |
| 15 | WISP1 | 5.56 |
| 16 | CTHRC1 | 5.55 |
| 17 | TENM3 | 5.34 |
| 18 | CLASRP | 5.30 |
| 19 | ARSI | 5.00 |
| 20 | ST3GAL6 | 4.89 |

| **Breast Invasive Carcinoma** | | |
| --- | --- | --- |
| **Ranking** | **Host Genes** | **Contribution (×10^­-3^)** |
| 1 | POLD4 | 5.80 |
| 2 | SEMA3B | 4.86 |
| 3 | INAFM1 | 4.10 |
| 4 | GMPS | 3.74 |
| 5 | FLT3LG | 3.62 |
| 6 | TAF2 | 3.16 |
| 7 | TNFSF12 | 3.16 |
| 8 | KLF2 | 3.13 |
| 9 | PTRF | 3.04 |
| 10 | RC3H2 | 2.93 |
| 11 | NUP155 | 2.80 |
| 12 | RPL7L1 | 2.78 |
| 13 | IGFBP6 | 2.73 |
| 14 | RAD23B | 2.70 |
| 15 | RCN2 | 2.70 |
| 16 | ZNF688 | 2.67 |
| 17 | RFWD3 | 2.61 |
| 18 | IFFO1 | 2.56 |
| 19 | XPOT | 2.55 |
| 20 | OPA1 | 2.53 |

| **Cervical Squamous Cell Carcinoma and Endocervical Adenocarcinoma** | | |
| --- | --- | --- |
| **Ranking** | **Host Genes** | **Contribution (×10^­-3^)** |
| 1 | CCDC85C | 6.16 |
| 2 | MAPK8IP3 | 6.14 |
| 3 | ZNF316 | 4.37 |
| 4 | EMCN | 3.66 |
| 5 | FAM219B | 3.45 |
| 6 | SMIM7 | 3.40 |
| 7 | TPGS1 | 3.24 |
| 8 | PLEC | 3.16 |
| 9 | RAI1 | 2.92 |
| 10 | TMEM131 | 2.89 |
| 11 | ARHGEF11 | 2.77 |
| 12 | FRG1 | 2.75 |
| 13 | CTU1 | 2.71 |
| 14 | UCHL3 | 2.69 |
| 15 | TMEM132C | 2.67 |
| 16 | COMMD9 | 2.63 |
| 17 | RP11-691N7.6 | 2.63 |
| 18 | SYF2 | 2.61 |
| 19 | DNAJC8 | 2.59 |
| 20 | OCLM | 2.58 |

| **Colon Adenocarcinoma** | | |
| --- | --- | --- |
| **Ranking** | **Host Genes** | **Contribution (×10^­-3^)** |
| 1 | PRR36 | 6.74 |
| 2 | TMED5 | 4.84 |
| 3 | RRNAD1 | 4.64 |
| 4 | PLXNB1 | 4.13 |
| 5 | CYHR1 | 3.49 |
| 6 | ERI1 | 3.35 |
| 7 | RGL2 | 3.25 |
| 8 | VPS37A | 3.23 |
| 9 | ZNF219 | 3.01 |
| 10 | CTSC | 2.97 |
| 11 | PAQR6 | 2.91 |
| 12 | UPK1A | 2.91 |
| 13 | EVX1 | 2.90 |
| 14 | USP33 | 2.87 |
| 15 | B2M | 2.82 |
| 16 | CSTL1 | 2.80 |
| 17 | TRIM11 | 2.77 |
| 18 | GOSR2 | 2.74 |
| 19 | WBP1 | 2.71 |
| 20 | FBXL8 | 2.69 |

| **Esophageal Aarcinoma** | | |
| --- | --- | --- |
| **Ranking** | **Host Genes** | **Contribution (×10^­-3^)** |
| 1 | FOXI3 | 10.32 |
| 2 | SIGIRR | 8.96 |
| 3 | TTBK2 | 8.93 |
| 4 | KCTD15 | 8.84 |
| 5 | B3GALNT2 | 7.88 |
| 6 | MARVELD1 | 7.11 |
| 7 | TMEM125 | 6.67 |
| 8 | SLC44A3 | 6.43 |
| 9 | CYP3A5 | 6.39 |
| 10 | CLDN23 | 5.91 |
| 11 | MMEL1 | 5.63 |
| 12 | CHODL | 5.54 |
| 13 | CRACR2B | 5.48 |
| 14 | FAM109A | 5.42 |
| 15 | TMPRSS2 | 5.35 |
| 16 | RP11-514O12.4 | 5.35 |
| 17 | EFCAB1 | 5.34 |
| 18 | NUP93 | 5.22 |
| 19 | C17orf53 | 5.07 |
| 20 | VIL1 | 4.96 |

| **Head and Neck Squamous Cell Carcinoma** | | |
| --- | --- | --- |
| **Ranking** | **Host Genes** | **Contribution (×10^­-3^)** |
| 1 | CAV2 | 6.18 |
| 2 | ARHGEF33 | 4.75 |
| 3 | TSC2 | 3.95 |
| 4 | PSMC1 | 3.63 |
| 5 | ARHGAP4 | 3.59 |
| 6 | GALNT3 | 3.48 |
| 7 | TNFRSF13B | 3.46 |
| 8 | ZNF322 | 3.45 |
| 9 | RAB18 | 3.44 |
| 10 | EFNB2 | 3.42 |
| 11 | TCP11 | 3.38 |
| 12 | LIPA | 3.30 |
| 13 | UPB1 | 3.28 |
| 14 | TYK2 | 3.13 |
| 15 | XPR1 | 3.12 |
| 16 | IKZF3 | 3.06 |
| 17 | ZNF541 | 3.05 |
| 18 | PDXK | 2.93 |
| 19 | CDKN2C | 2.92 |
| 20 | SMC1B | 2.89 |

| **Kidney Chromophobe** | | |
| --- | --- | --- |
| **Ranking** | **Host Genes** | **Contribution (×10^­-3^)** |
| 1 | SPATA5 | 25.53 |
| 2 | NUP155 | 22.05 |
| 3 | UTP15 | 15.92 |
| 4 | RINT1 | 14.39 |
| 5 | ATRX | 14.23 |
| 6 | RFC5 | 13.22 |
| 7 | RSBN1 | 12.99 |
| 8 | CUL5 | 12.46 |
| 9 | PRPF8 | 11.76 |
| 10 | GPM6B | 10.13 |
| 11 | PPWD1 | 9.00 |
| 12 | ZW10 | 8.26 |
| 13 | NAT10 | 8.24 |
| 14 | NUP107 | 8.22 |
| 15 | PCNP | 8.10 |
| 16 | ZNF107 | 8.09 |
| 17 | ZCRB1 | 7.78 |
| 18 | UBXN7 | 7.74 |
| 19 | FBXW11 | 7.69 |
| 20 | HAUS6 | 7.68 |

| **Kidney Renal Clear Cell Carcinoma** | | |
| --- | --- | --- |
| **Ranking** | **Host Genes** | **Contribution (×10^­-3^)** |
| 1 | PPAP2B | 17.77 |
| 2 | TAF10 | 13.33 |
| 3 | OSBPL1A | 11.65 |
| 4 | FBXL3 | 10.74 |
| 5 | ITGA6 | 10.23 |
| 6 | SUCLA2 | 9.81 |
| 7 | SEPP1 | 9.00 |
| 8 | NPEPL1 | 8.70 |
| 9 | LRBA | 8.33 |
| 10 | FBF1 | 8.29 |
| 11 | CAT | 8.17 |
| 12 | PPP6C | 7.65 |
| 13 | TCIRG1 | 7.38 |
| 14 | TSPYL1 | 7.16 |
| 15 | HDHD2 | 7.11 |
| 16 | SPC24 | 6.68 |
| 17 | ACADSB | 6.24 |
| 18 | ENPP5 | 5.79 |
| 19 | ACADM | 5.72 |
| 20 | HN1 | 5.54 |

| **Kidney Renal Papillary Cell Carcinoma** | | |
| --- | --- | --- |
| **Ranking** | **Host Genes** | **Contribution (×10^­-3^)** |
| 1 | TOP2A | 17.42 |
| 2 | BLM | 15.00 |
| 3 | CCNB1 | 12.71 |
| 4 | NEK2 | 12.46 |
| 5 | SKA1 | 11.98 |
| 6 | MCM6 | 11.95 |
| 7 | LMNB1 | 11.52 |
| 8 | ARHGAP11A | 10.34 |
| 9 | CDC25C | 10.00 |
| 10 | MND1 | 9.78 |
| 11 | TTK | 9.76 |
| 12 | KIF11 | 9.11 |
| 13 | KIF18A | 8.70 |
| 14 | RSL1D1 | 8.41 |
| 15 | SPATA18 | 7.51 |
| 16 | CKAP2L | 7.43 |
| 17 | CCNB2 | 7.41 |
| 18 | DEPDC1B | 7.13 |
| 19 | RACGAP1 | 7.08 |
| 20 | TPX2 | 6.95 |

| **Brain Lower Grade Glioma** | | |
| --- | --- | --- |
| **Ranking** | **Host Genes** | **Contribution (×10^­-3^)** |
| 1 | GAS2L3 | 20.97 |
| 2 | SLC30A7 | 19.37 |
| 3 | EVC2 | 15.42 |
| 4 | ATP6V1G2 | 13.97 |
| 5 | SMC4 | 13.77 |
| 6 | FNDC3B | 13.04 |
| 7 | PLAT | 12.39 |
| 8 | EFEMP2 | 11.74 |
| 9 | STIL | 11.38 |
| 10 | ADAM12 | 11.29 |
| 11 | FAIM2 | 11.20 |
| 12 | ZNF217 | 10.18 |
| 13 | RBMS1 | 10.02 |
| 14 | ZNF28 | 8.02 |
| 15 | PRLHR | 7.90 |
| 16 | KDELR2 | 7.84 |
| 17 | PLP2 | 7.55 |
| 18 | ZWILCH | 7.06 |
| 19 | PELI3 | 7.01 |
| 20 | CRY2 | 6.91 |

| **Liver Hepatocellular Carcinoma** | | |
| --- | --- | --- |
| **Ranking** | **Host Genes** | **Contribution (×10^­-3^)** |
| 1 | NEU3 | 3.27 |
| 2 | NFASC | 2.52 |
| 3 | ETS2 | 2.48 |
| 4 | NDRG1 | 2.46 |
| 5 | HAVCR1 | 2.43 |
| 6 | DEK | 2.35 |
| 7 | BLOC1S6 | 2.30 |
| 8 | FAM206A | 2.29 |
| 9 | IL33 | 2.18 |
| 10 | EFTUD1 | 2.12 |
| 11 | ATG101 | 2.11 |
| 12 | EDIL3 | 2.09 |
| 13 | OSBPL3 | 1.91 |
| 14 | NUPL1 | 1.90 |
| 15 | NDUFB7 | 1.88 |
| 16 | SREK1 | 1.86 |
| 17 | LYPD8 | 1.82 |
| 18 | PHF6 | 1.82 |
| 19 | GNAI2 | 1.75 |
| 20 | IFIH1 | 1.72 |

| **Lung Squamous Cell Carcinoma** | | |
| --- | --- | --- |
| **Ranking** | **Host Genes** | **Contribution (×10^­-3^)** |
| 1 | ZNF628 | 3.83 |
| 2 | GPKOW | 2.44 |
| 3 | TIA1 | 2.41 |
| 4 | EMC6 | 2.37 |
| 5 | PSMG1 | 2.21 |
| 6 | PNN | 2.15 |
| 7 | AL158801.1 | 2.05 |
| 8 | SLC5A3 | 2.05 |
| 9 | NCKAP1 | 2.02 |
| 10 | SERPIND1 | 2.01 |
| 11 | LRRC48 | 1.94 |
| 12 | ZNF443 | 1.85 |
| 13 | SCRN3 | 1.83 |
| 14 | RAB33A | 1.82 |
| 15 | TXN | 1.70 |
| 16 | MAP10 | 1.69 |
| 17 | FHL3 | 1.63 |
| 18 | TNFRSF10B | 1.58 |
| 19 | IER5L | 1.58 |
| 20 | C14orf80 | 1.58 |

| **Pancreatic Adenocarcinoma** | | |
| --- | --- | --- |
| **Ranking** | **Host Genes** | **Contribution (×10^­-3^)** |
| 1 | ANXA6 | 14.71 |
| 2 | NLRP1 | 14.67 |
| 3 | GRHL2 | 12.92 |
| 4 | PPM1K | 11.93 |
| 5 | PPP1R3E | 11.79 |
| 6 | ST3GAL3 | 10.78 |
| 7 | PRCD | 10.63 |
| 8 | NRG2 | 8.93 |
| 9 | FERMT1 | 8.80 |
| 10 | MROH8 | 8.47 |
| 11 | CD99L2 | 8.26 |
| 12 | MMD2 | 8.25 |
| 13 | TBC1D10C | 7.65 |
| 14 | PKD1 | 7.40 |
| 15 | F11R | 7.19 |
| 16 | TERF2IP | 7.06 |
| 17 | OTX1 | 6.85 |
| 18 | POU6F1 | 6.64 |
| 19 | PAIP2 | 6.64 |
| 20 | CRIP3 | 6.57 |

| **Rectum Adenocarcinoma** | | |
| --- | --- | --- |
| **Ranking** | **Host Genes** | **Contribution (×10^­-3^)** |
| 1 | PRRC2B | 10.26 |
| 2 | CNOT1 | 10.14 |
| 3 | TRAPPC10 | 9.97 |
| 4 | CCNF | 9.65 |
| 5 | MIEF1 | 9.58 |
| 6 | HNRNPM | 9.36 |
| 7 | PNP | 7.56 |
| 8 | PARD3 | 7.14 |
| 9 | MAFG | 6.57 |
| 10 | FASN | 6.55 |
| 11 | CCNB1 | 6.09 |
| 12 | IARS | 5.94 |
| 13 | MTM1 | 5.78 |
| 14 | IL7R | 5.66 |
| 15 | MYH9 | 5.44 |
| 16 | AGAP1 | 5.16 |
| 17 | ZNF318 | 4.80 |
| 18 | NCAPG2 | 4.79 |
| 19 | JOSD1 | 4.75 |
| 20 | PVR | 4.65 |

| **Sarcoma** | | |
| --- | --- | --- |
| **Ranking** | **Host Genes** | **Contribution (×10^­-3^)** |
| 1 | PTK2B | 8.39 |
| 2 | NDC1 | 8.31 |
| 3 | LMNB2 | 7.98 |
| 4 | CCDC69 | 6.91 |
| 5 | GLO1 | 6.45 |
| 6 | FAM171A2 | 6.41 |
| 7 | PCED1B | 5.70 |
| 8 | EXOSC10 | 5.33 |
| 9 | CD40LG | 5.32 |
| 10 | YIF1B | 5.17 |
| 11 | BLNK | 4.90 |
| 12 | TNFRSF14 | 4.75 |
| 13 | GRAP2 | 4.70 |
| 14 | DPYS | 4.64 |
| 15 | SAPCD2 | 4.56 |
| 16 | SH3BP4 | 4.55 |
| 17 | HLA-F | 4.46 |
| 18 | ANKS6 | 4.35 |
| 19 | DFFA | 4.31 |
| 20 | A2M | 4.14 |

| **Skin Cutaneous Melanoma** | | |
| --- | --- | --- |
| **Ranking** | **Host Genes** | **Contribution (×10^­-3^)** |
| 1 | NR2F6 | 13.09 |
| 2 | CCL3 | 10.11 |
| 3 | CYP19A1 | 8.11 |
| 4 | APOOL | 7.49 |
| 5 | NRTN | 7.29 |
| 6 | NDRG2 | 6.99 |
| 7 | JUND | 6.55 |
| 8 | HIST2H2BF | 6.09 |
| 9 | MAGI1 | 5.54 |
| 10 | PKP4 | 5.18 |
| 11 | NSMF | 5.16 |
| 12 | CADPS2 | 5.05 |
| 13 | HIST1H1E | 4.99 |
| 14 | BOD1 | 4.89 |
| 15 | NMB | 4.86 |
| 16 | FAM173A | 4.86 |
| 17 | INO80 | 4.84 |
| 18 | LSMEM2 | 4.76 |
| 19 | NTNG2 | 4.64 |
| 20 | SLC16A2 | 4.62 |

| **Thymoma** | | |
| --- | --- | --- |
| **Ranking** | **Host Genes** | **Contribution (×10^­-3^)** |
| 1 | HMGA1 | 10.60 |
| 2 | TARBP2 | 9.89 |
| 3 | CHRNA5 | 8.58 |
| 4 | MEN1 | 7.64 |
| 5 | DNASE1L1 | 7.47 |
| 6 | LPAR6 | 7.09 |
| 7 | TAF15 | 7.08 |
| 8 | ARHGEF37 | 6.02 |
| 9 | NUDT5 | 5.93 |
| 10 | FAM63A | 5.91 |
| 11 | DTL | 5.58 |
| 12 | KLHL22 | 5.54 |
| 13 | NBPF9 | 5.47 |
| 14 | POLR1D | 5.45 |
| 15 | SLC8B1 | 5.42 |
| 16 | ADAM28 | 5.35 |
| 17 | IQGAP2 | 5.27 |
| 18 | RNF19B | 5.22 |
| 19 | PHYH | 5.21 |
| 20 | SYNGR1 | 5.18 |

| **Uterine Corpus Endometrial Carcinoma** | | |
| --- | --- | --- |
| **Ranking** | **Host Genes** | **Contribution (×10^­-3^)** |
| 1 | IHH | 15.36 |
| 2 | SPDEF | 14.68 |
| 3 | PPAP2C | 13.26 |
| 4 | GID8 | 10.77 |
| 5 | DDX27 | 10.01 |
| 6 | PDCL2 | 8.11 |
| 7 | TRPM4 | 7.64 |
| 8 | DLL3 | 6.59 |
| 9 | SYT13 | 6.12 |
| 10 | EPHB2 | 6.07 |
| 11 | TMEM41A | 6.02 |
| 12 | URI1 | 5.86 |
| 13 | KCNK6 | 5.80 |
| 14 | ELOVL3 | 5.69 |
| 15 | EEF2 | 5.60 |
| 16 | HPDL | 5.45 |
| 17 | TTI1 | 4.97 |
| 18 | L1CAM | 4.90 |
| 19 | OSGIN1 | 4.75 |
| 20 | PBX2 | 4.40 |

| **Uveal Melanoma** | | |
| --- | --- | --- |
| **Ranking** | **Host Genes** | **Contribution (×10^­-3^)** |
| 1 | DLL4 | 32.38 |
| 2 | NR6A1 | 27.03 |
| 3 | BRK1 | 18.04 |
| 4 | MTMR14 | 17.95 |
| 5 | TSGA10IP | 17.06 |
| 6 | SSUH2 | 16.59 |
| 7 | RNF43 | 16.42 |
| 8 | AIFM2 | 16.17 |
| 9 | SLC25A38 | 16.06 |
| 10 | TMEM41A | 14.93 |
| 11 | GJC3 | 14.50 |
| 12 | PLEKHG6 | 14.40 |
| 13 | DALRD3 | 9.51 |
| 14 | SLC25A26 | 9.49 |
| 15 | SLC41A3 | 9.48 |
| 16 | RPL32 | 9.46 |
| 17 | GPR156 | 9.03 |
| 18 | HTR2B | 9.01 |
| 19 | FERMT3 | 9.01 |
| 20 | MORC2 | 9.01 |
